# Nanoscale architecture of the axon initial segment reveals an organized and robust scaffold

**DOI:** 10.1101/022962

**Authors:** Christophe Leterrier, Jean Potier, Ghislaine Caillol, Claire Debarnot, Fanny Rueda Boroni, Bénédicte Dargent

**Affiliations:** Aix Marseille Université, CNRS, CRN2M UMR7286, 13344 cedex 15, Marseille, France

**Keywords:** axon initial segment, cytoskeleton, ankyrin G, super-resolution microscopy

## Abstract

The Axon Initial Segment [AIS], located within the first 30 μm of the axon, has two essential roles in generating action potentials and maintaining axonal identity. AIS assembly depends on a ßIV-spectrin / ankyrin G scaffold, but its macromolecular arrangement is not well understood. Here we quantitatively determined the AIS nanoscale architecture using STochastic Optical Reconstruction Microscopy [STORM]. First we directly demonstrate that the 190-nm periodicity of the AIS submembrane lattice results from longitudinal, head-to-head ßIV-spectrin molecules connecting actin rings. Using multicolor 3D-STORM, we resolve the nanoscale organization of ankyrin G: its aminoterminus associates with the submembrane lattice, whereas the carboxyterminus radially extends (~32 nm on average) toward the cytosol. This AIS nano-architecture is highly resistant to cytoskeletal perturbations, advocating its role in structural stabilization. Our findings provide a comprehensive view of the AIS molecular architecture, and will help understanding the crucial physiological functions of this compartment.

## Introduction

The directional flow of information in the brain is ensured by the cellular asymmetry of the neuron: the neuronal cell body receives synaptic inputs, and the axon propagates the action potential to downstream neurons. Neuronal asymmetry is maintained for years or even decades by a combination of passive (barriers) and active (directed traffic) processes, but the underpinning mechanisms remain largely unknown (Kapitein and Hoogenraad, 2011). Located along the first 20 to 40 μm of the axon, the Axon Initial Segment [AIS] materializes the separation between the cell body and the axon. This location allows for the two main cellular functions of the AIS: the initiation of action potentials, and the maintenance of axonal identity (Leterrier and Dargent, 2014).

The AIS ensures the proper generation of action potentials by concentrating voltage-gated sodium (Nav) and potassium (Kv7) channels at its surface. Together with specific cell adhesion molecules [CAMs] such as 186-kDa neurofascin, these channels are anchored by an interaction with the specialized AIS scaffold protein ankyrin G [ankG]. AnkG also binds to a submembrane complex of ßIV-spectrin and actin (for reviews see Bennett and Lorenzo, 2013; Grubb and Burrone, 2010a; Leterrier and Dargent, 2014; Normand and Rasband, 2015). Furthermore, ankG links the AIS scaffold to microtubule fascicles, via an interaction with end-binding proteins EB1 and EB3 (Leterrier et al., 2011). AnkG is the AIS master organizer: depletion of ankG results in the absence or disassembly of the whole AIS complex (Hedstrom et al., 2008; 2007; S. M. Jenkins and Bennett, 2001). In ankG-depleted neurons that lack an AIS, somatodendritic proteins progressively invade the proximal axon, resulting in the disappearance of microtubules fascicles, and the formation of ectopic post-synapses (Hedstrom et al., 2008; Sobotzik et al., 2009): this demonstrated the role of the AIS for the maintenance of axonal identity.

Two proposed cellular processes contribute to this maintenance of axonal identity: a surface diffusion barrier that restricts the mobility of membrane proteins and lipids (Nakada et al., 2003; Winckler et al., 1999), and a traffic filter that could regulate intracellular diffusion and vesicular transport (Song et al., 2009). The underlying mechanisms are still mysterious, in particular the nature of the intracellular filter (Petersen et al., 2014; Watanabe et al., 2012). Our understanding of these processes depends on a better knowledge of the AIS architecture down to the molecular level. Super-resolution microscopy now allows to observe macromolecular complexes in situ with a resolution down to a few tens of nanometers (Maglione and Sigrist, 2013), and has recently started to uncover the nanoscale organization of the AIS and axon. STochastic Optical Reconstruction Microscopy [STORM] has revealed that axonal actin is organized as submembrane rings periodically spaced every 190 nm (Xu et al., 2013), a result recently confirmed in living cells (D’Este et al., 2015). In the AIS, a periodic arrangement of actin, ßIV-spectrin and ankG is also detected (Zhong et al., 2014). However, the relative arrangement of AIS components resulting in this regular organization has not been directly addressed. Furthermore, although electron microscopy recently resolved individual ankG proteins as 150 nm-long rods (Jones et al., 2014), the nanoscale organization of this key protein within the AIS scaffold is still elusive. Notably, a connection between the plasma membrane components and intracellular structures by large isoforms of ankG has been proposed (Davis et al., 1996; Leterrier and Dargent, 2014) but remains hypothetical.

We set about quantitatively resolving the three-dimensional architecture of the AIS at the nanoscale level, in particular the arrangement of the master scaffold protein ankG. Multicolor 2D- and 3D-STORM of endogenous epitopes coupled to extensive quantification procedures allowed us to uncover the intricate ordering of the AIS scaffold. We directly demonstrate that the submembrane periodic lattice is composed of longitudinal head-to-head ßIV-spectrin molecules connecting submembrane actin bands along the AIS. AnkG aminoterminal side is associated with this lattice, and its carboxyterminal side extends away from the plasma membrane, ~35 nm deeper in the cytoplasm. This could allow its interaction with peripheral microtubules, providing a structural basis for the regulation of vesicular entry into the axon. Finally, our observations reveal an unexpected robustness of the AIS scaffold: its ordered arrangement is resistant to pharmacological perturbations of the actin or microtubule cytoskeleton, and the radial extent of ankG even resists partial disassembly of the AIS induced by elevated intracellular calcium. This suggests that the nanoscale organization of AIS components is necessary for its integrity, and this structural robustness may support the AIS role as a gateway to the axon.

## Results

### Actin and longitudinal head-to-head ßIV-spectrin molecules form a periodic submembrane complex at the AIS

To characterize the AIS nano-architecture, we first focused on the organization of the submembrane actin/**ß**IV-spectrin complex. At the diffraction-limited level, actin was present but not concentrated in the AIS of mature neurons (15 to 21 days in vitro), in contrast to **ß**IV-spectrin and Nav channels (Figure 1A). We used STORM that provided a ~17 nm lateral localization precision (see Figure S1A-C and Extended Experimental Procedures) to obtain images of the nanoscale actin organization. 2D-STORM images revealed the periodic arrangement of AIS actin as regular bands (Figure 1B-C), as described previously (D’Este et al., 2015; Xu et al., 2013). Line profiles obtained on STORM images exhibited intensity peaks with a regular spacing of ~190 nm (Figure 1D). We quantified the labeling periodicity by fitting sinusoids on 1 **μ**m-long intensity profiles, and fitting of the resulting spacing values histogram with a Gaussian curve (Figure S1D-G). The mean spacing and its spread were measured at 188 ± 8 nm for actin labeling along the AIS (Table S1). We next assessed the labeling obtained with anti **ß**IV-spectrin antibodies against either its aminoterminus [NT], which binds actin, or its specific domain [SD], close to the carboxyterminus. Both **ß**IV-spectrin NT (Figure 1E-F) and **ß**IV-spectrin SD (Figure 1H-I) exhibited a strikingly periodic labeling pattern, and we measured the spacing distribution at 191 ± 7 nm for **ß**IV-spectrin NT (Figure 1G), and 188 ± 8 nm for **ßI**V-spectrin SD (Figure 1J), in close agreement with values reported previously (Xu et al., 2013; Zhong et al., 2014). As observed for actin by us and others (D’Este et al., 2015), AIS of large diameter showed a more complex ßIV-spectrin arrangement, with apposed stretches of bands resembling an herringbone parquet flooring (see Figure 2C for an example).

**Figure 1. (previous page).**
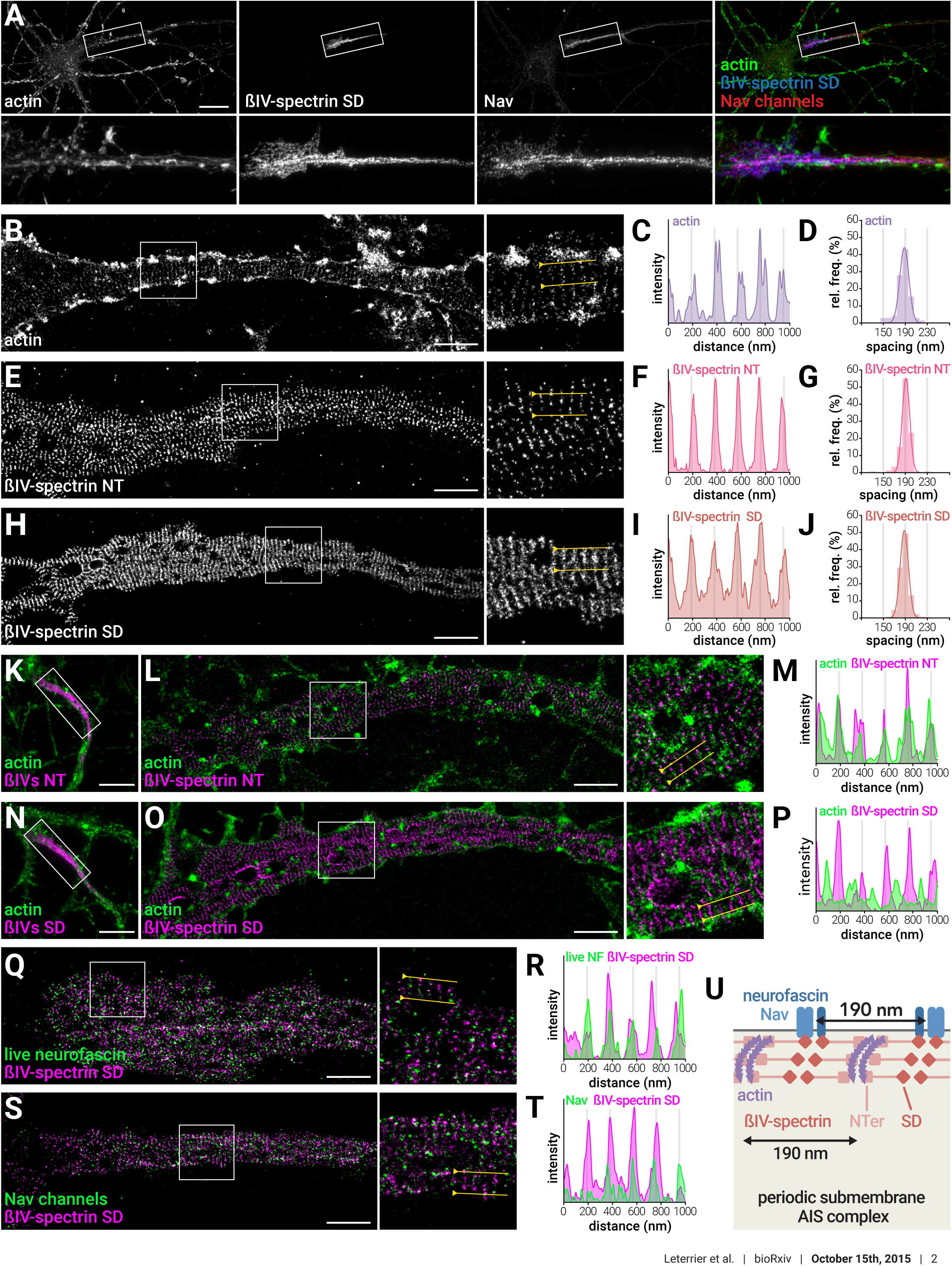
Actin and longitudinal head-to-head ßIV-spectrin molecules form a periodic submembrane complex at the AIS. (A) Epifluorescence image of a neuron labeled for actin (phalloidin, green), ßIV-spectrin SD (blue) and Nav channels (red). Scale bar, 20 μm. (B) STORM image of an AIS labeled for actin. Scale bars for (B, E, H, L, O, Q, S) are 2 μm. (C) Intensity profile along the yellow line (gray lines are 190 nm apart). (D) Histogram of the spacing values (n=72 profiles from N=3 independent experiments). (E-G) Same as (B-D), for an AIS labeled for ßIV-spectrin NT (histogram, n=183 N=3). (H-J) Same as (B-D), for an AIS labeled for ßIV-spectrin SD (histogram, n=334 N=11). (K) Epifluorescence image of a neuron labeled for actin (green) and ßIV-spectrin NT (magenta). Scale bars for (K, N) are 10 μm. (L) 2-color STORM image of the AIS shown in (K); (M) corresponding intensity profile. (N-P) Same as (K-M), for a neuron labeled (green) and ßIV-spectrin SD (magenta). (Q) 2-color STORM image of an AIS labeled live for neurofascin (NF, green) and ßIV-spectrin SD (magenta); (R) corresponding intensity profile. (S-T) Same as (Q-R), for a neuron labeled for Nav channels (green) and ßIV-spectrin SD (magenta). (U) Structural model of the submembrane AIS complex. See also Figure S1 and S2.

**Figure 2.**
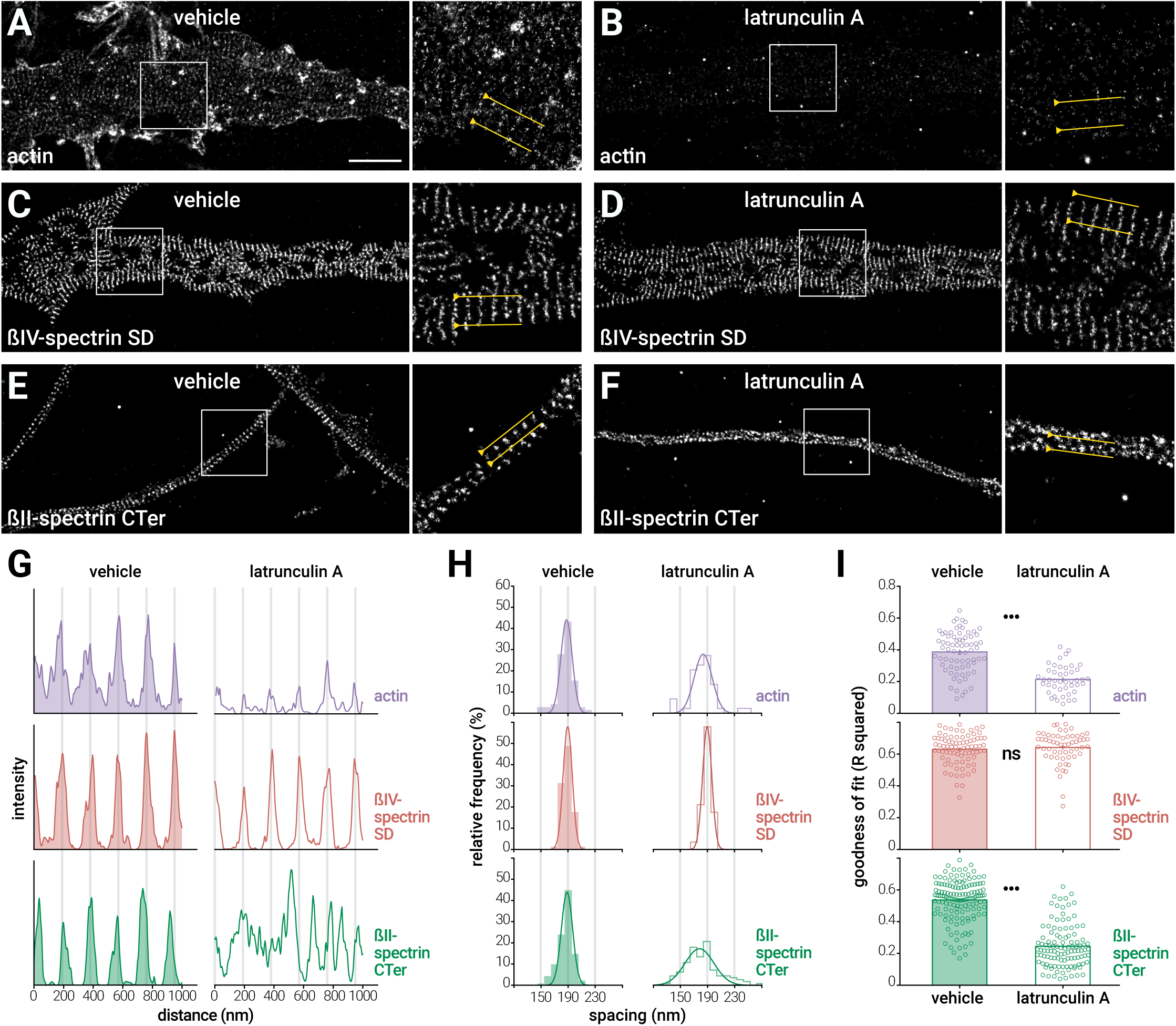
The AIS submembrane lattice is resistant to actin perturbation. (A-F) STORM images of neurons treated with vehicle (DMSO 0.1%, 1h) (A, C, E) or latrunculin A (latA, 5μM, 1h) (B, D, F), fixed and labeled for actin (AIS: A, B), ßIV-spectrin SD (AIS: C, D) or ßII-spectrin (distal axons: E, F). Scale bars for (A-F) are 2 μm. (G) Intensity profiles along the yellow lines in (A-F) for the actin (purple), ßIV-spectrin SD (red) and ßII-spectrin CTer (green) labeling. Intensities have been processed identically between the vehicle (left) and latA (right) treated conditions. (H) Histograms of the spacing values for each labeling (n=44-143 profiles, N=3–4). (I) Goodness of sinusoid fit (R squared) for each labeling (n=44-143, N=3–4). See also Figure S3.

We next directly assessed the relative organization of actin and ßIV-spectrin using two-color STORM of actin together with ßIV-spectrin NT or SD (Figure 1K-P). Although diffraction-limited level images were similar (Figure 1K and 1N), on STORM images the periodic ßIV-spectrin NT bands colocalized along actin bands (Figure 1L-M), whereas the ßIV-spectrin SD bands were alternating with actin bands (Figure 1O-P). This differential position of ßIV-spectrin ends relative to actin bands directly demonstrates the proposed model of head-to-head ßIV-spectrin molecules connecting actin bands along the AIS (Figure 1U). Finally, we visualized the nanoscale distribution of two AIS membrane proteins shown to localize along the periodic submembrane complex, 186-kDa neurofascin (D’Este et al., 2015) and Nav channels (Xu et al., 2013). Single-color STORM images revealed a clustered organization of neurofascin and Nav channels, and we could detect the presence of some periodicity in the cluster arrangement (Figure S2A-B and S2D-E), although it was quite low, resulting in flattened spacing histograms (190 ± 16 nm and 188 ± 18 nm for neurofascin and Nav channels, respectively). Two-color STORM revealed that the neurofascin and Nav clusters localized along ßIV-spectrin SD bands, confirming their underlying periodicity (Figure 1Q-T). Overall, our data validate and extend previous reports (D’Este et al., 2015; Xu et al., 2013; Zhong et al., 2014), and provides a comprehensive view for the molecular structure of the AIS submembrane complex (Figure 1U).

### The AIS submembrane lattice is resistant to actin perturbation

The AIS scaffold is remarkably stable over time (Hedstrom et al., 2008). We wondered if the ßIV-spectrin periodic lattice would nonetheless depend on integrity of the actin structure. To test this hypothesis, we assessed the ßIV-spectrin nanoscale distribution after actin depolymerization by latrunculin A [latA]. Treatment with 5 μM latA for one hour disassembled actin filaments in all compartments, with a near disappearance of phalloidin labeling on diffraction-limited images and an 85% drop of labeling intensity at the AIS (Figure S3A-C). Notably, this short term treatment did not affect the concentration of AIS components or the AIS length (Figure S3C-D).

STORM images revealed the disorganization of actin in the AIS, although a faint remnant of actin periodicity could still be detected, suggesting a selective resistance to depolymerization (Figure 2A-B & 2G). Strikingly, the periodic pattern of the ßIV-spectrin SD was unaffected by latA treatment (Figure 2C-D & 2G). In the control condition, actin and ßIV-spectrin in the AIS exhibited an identical spacing of 188 ± 8 nm (Figure 2H). After latA treatment, the spread of the spacing remained stable for ßIV-spectrin (190 ± 7 nm). Actin showed a significant drop in the goodness of sinusoid fit after latA treatment, indicating a loss of periodicity (from 0.39 ± 0.02 to 0.21 ± 0.01, mean ± SEM), but the goodness of fit for ßIV-spectrin remained high after actin depolymerization (0.63 ± 0.01 for vehicle, 0.64 ± 0.01 for latA, Figure 2I). Thus, periodic actin seems partially resistant to latA, and ßIV-spectrin periodic organization in the AIS is resistant to actin perturbation.

To determine if this robustness is specific to the AIS, we assessed the effect of actin disruption on the distal axon actin / spectrin complex, which forms a similar periodic structure with ßII-spectrin instead of the AIS-specific ßIV-spectrin (Xu et al., 2013). Using an antibody against the ßII-spectrin carboxyterminus (CTer), we indeed found a highly periodic distribution along distal axons, with a regular spacing of 188 ± 8 nm (Figure 2E & 2G). ßII-spectrin was still clustered after LatA treatment, but the regular spacing was disorganized, with a rising spacing spread (190 ± 22 nm, Figure 2H), and a drop of the sinusoid fit R squared from 0.54 ± 0.01 to 0.25 ± 0.01 (Figure 2I), confirming previous results (Xu et al., 2013; Zhong et al., 2014). In conclusion, we found actin to be partially resistant to depolymerization in the AIS. In contrast to the actin / ßII-spectrin complex found in the distal axon, the ßIV-spectrin periodic lattice is completely resistant to actin perturbation by latA.

### The ankG spectrin-binding domain associates with the periodic lattice, but its carboxyterminal part is not periodically arranged

What makes the actin / ßIV-spectrin lattice in the AIS more resistant than the ßII-spectrin/actin complex along the distal axon? Stabilization is likely due to the AIS master organizer ankG that recruits and maintains ßIV-spectrin at the AIS (Yang et al., 2007). AnkG present at the AIS is a large 480-kDa protein, with a shorter 270-kDa isoform being also expressed (Bennett and Lorenzo, 2013; P. M. Jenkins et al., 2015). To characterize the nanoscale organization of ankG by STORM, we used antibodies directed against distinct domains of the protein (Figure 3A). A polyclonal antibody directed against a peptide epitope in the serine-rich domain [SR] was previously described (Bréchet et al., 2008), and we mapped the target domains of two monoclonal antibodies (anti-ankG clone 106/65 and 106/36) to the spectrin-binding [SB] and carboxyterminal [CTer] domains, respectively (Figure S4A). We also used a tail480 antibody recognizing the distal part of the tail specific to 480-kDa ankG, close to the CTer (Figure S4B). These antibodies labeled the AIS of hippocampal neurons in a similar fashion at the diffraction-limited level (Figure 3B-C).

**Figure 3.**
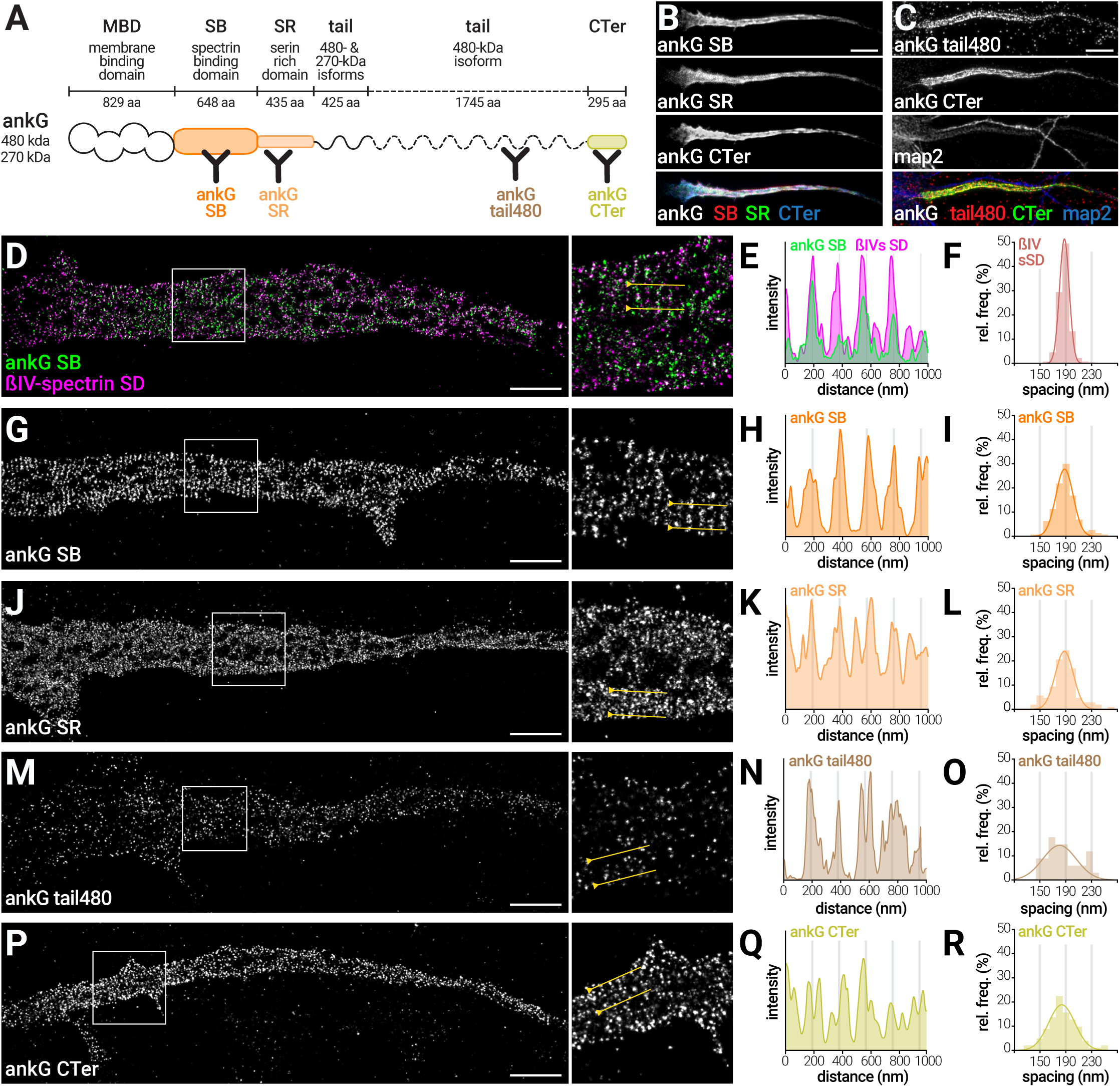
The ankG spectrin-binding domain associates with the periodic lattice, but its carboxyterminal part is not periodically arranged. (A) Cartoon of the ankG domains, with the target domains of the four anti-ankG antibodies used. (B) Epifluorescence image showing the AIS of a neuron labeled for ankG SB (red), ankG SR (green), and ankG CTer (blue). (C) Epifluorescence image showing the AIS of a neuron labeled for ankG tail480 (red), ankG CTer (red), and map2 (blue). Scale bars on (B, C) are 5 μm. (D) STORM image of an AIS labeled for ankG SB (green) and ßIV-spectrin SD (magenta). Scale bars for (D, G, J, M, P) are 2 μm. (E) Intensity profiles for each channel along the yellow line. (F) Histogram of the spacing values for the ßIV-spectrin SD labeling (same data as in J). (G) STORM image of an AIS labeled for ankG SB; (H) corresponding intensity profile; (I) histogram of spacing values (n=161 profiles, N =6). (J-L) Same as (G-I) for an AIS labeled for ankG SR (histogram, n=107 N =3). (M-O) Same as (G-I) for an AIS labeled for ankG tail480 (histogram, n=34 N =2). (P-R) Same as (G-I) for an AIS labeled for ankG CTer (histogram, n=102 N =4). See also Figure S4.

At the nanoscale level, we first localized the domain of ankG that interacts with ßIV-spectrin (the SB domain) together with ßIV-spectrin SD. Two-color STORM images showed that the ankG SB clusters localized along ßIV-spectrin bands, resulting in correlated line profiles (Figure 3D-E). We next compared the periodicity of labeling for four different ankG domains: the SR domain (localized adjacent to the SB domain), the distal tail domain (tail480 antibody) or the CTer domain (formed by the last 300 residues of the protein, Figure 3A). On single color STORM images, the regular spacing of ankG SB clusters was clearly detected (Figure 3G-H), as reflected by periodicity measurements (spacing 188 ± 13 nm, Figure 3I and Table S1). Periodicity was similar for the adjacent SR domain (Figure 3J-K), with a spacing of 188 ± 14 nm (Figure 3L). Both the distal tail and the CTer domain exhibited a clustered distribution, but periodic patterns were difficult to discern (Figure 3M-N and 3P-Q), and the spacing histograms exhibited a significantly higher spread (182 ± 27 and 184 ± 20 nm for the distal tail and CTer, respectively, Figure 3O-R). Accordingly, the goodness of sinusoid fit (R squared) dropped from 0.34 ± 0.01 for the ankG SB domain, to 0.26 ± 0.01 for the SR, 0.20 ± 0.01 for the distal tail and 0.22 ± 0.01 for the CTer domain (Table S1). This gradual loss of periodic distribution between the ankG SB and CTer domains validates and extends a recent observation (Zhong et al., 2014). In conclusion, ankG exhibits a periodic arrangement of its aminoterminal side that interacts with ßIV-spectrin, but downstream domains, closer to its carboxyterminus, progressively loose this periodicity and become more disordered.

### The ankG CTer domain extends radially below the submembrane lattice

Why does the carboxyterminal part of ankG show a disorganized nanoscale distribution? We hypothesized it could extend away from the periodic submembrane lattice, deeper in the axoplasm where it could reach intracellular partners (see Figure 4O). To test this, we resolved the transverse organization of the AIS using 3D-STORM, which attained an axial localization precision of ~33 nm (see Figure S5A-C and Extended Experimental Procedures). To map the radial position of AIS components, we performed 2-color 3D-STORM with different epitopes co-labeled with ßIV-spectrin SD as a reference (Figure 4A-J). We first validated our method by imaging intracellular microtubules together with submembrane ßIV-spectrin SD (Figure 4A). Although *α*-tubulin labeling was not resolved as continuous microtubules due to the fixation protocol used in these experiments (compare with Figure 5G), intracellular labeling was nevertheless clear on transverse sections and radial profiles, with the ßIV-spectrin lattice encasing the *α*-tubulin labeling (Figure 4A-B). Next, we assessed the radial position of AIS membrane proteins: neurofascin using an antibody against an extracellular epitope (Figure 4C-D), and Nav channels using an antibody against an intracellular loop (Figure 4 E-F). On YZ transverse sections, both neurofascin and Nav channels were found at the periphery of the ßIV-spectrin submembrane lattice (Figure 4C and 4E), as confirmed by line profiles across transverse sections (Figure 4D and 4F).

**Figure 4.**
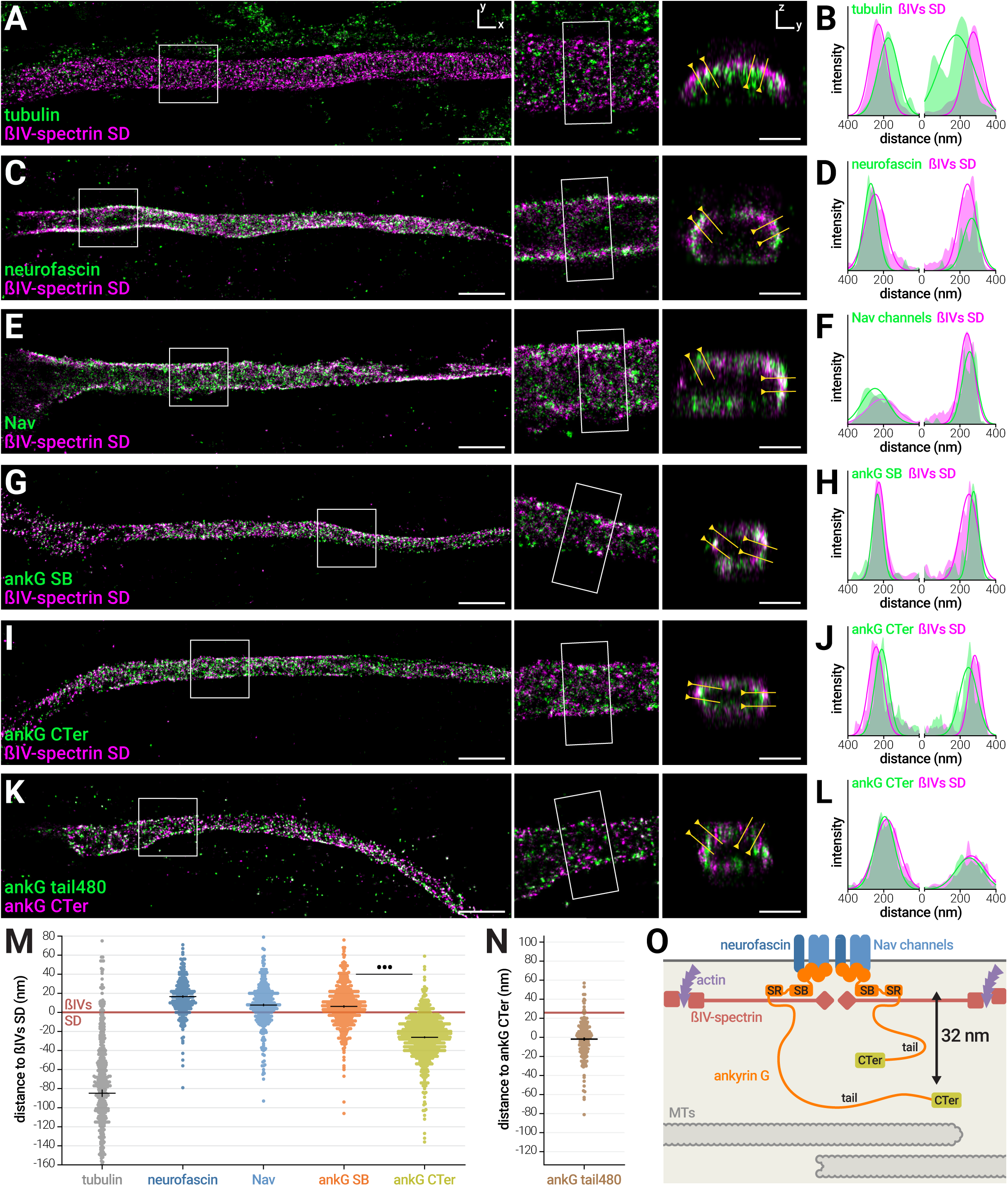
The ankG CTer domain extends radially below the submembrane lattice. (A) 3D-STORM image of an AIS labeled for α-tubulin (green) and ßIV-spectrin SD (magenta). (B) Intensity profiles for each channel (filled curves) and Gaussian fits used to calculate radial distances. For (A, C, E, G, I, K) profiles are taken along the yellow line on the corresponding transverse section shown, scale bars are 2 μm for XY image, 500 nm for YZ section. (C-D) Same as (A-B), for an AIS labeled for neurofascin (green) and ßIV-spectrin SD (magenta). (E-F) Same as (A-B), for an AIS labeled for Nav channels (green) and ßIV-spectrin SD (magenta). (G-H) Same as (A-B), for an AIS labeled for ankG SB (green) and ßIV-spectrin SD (magenta). (I-J) Same as (A-B), for an AIS labeled for ankG CTer (green) and ßIV-spectrin SD (magenta). (K-L) Same as (A-B), for an AIS labeled for ankG tail480 (green) and ankG CTer (magenta). (M) Radial distance to ßIV-spectrin SD for the α-tubulin, neurofascin, Nav, ankG SB, and ankG CTer labeling. Red line is ßIV-spectrin SD reference at 0 nm (n=284-832 profiles, N=2-9). (N) Radial distance between the ankG CTer and the ankG tail480 labeling (shifted Y scale to align with the ßIV-spectrin SD reference in (K), n=168, N=2). (O) Structural model of the AIS radial organization. See also Figure S5.

We then assessed the transverse arrangement of ankG by imaging the two domains on opposite side of the protein (SB and CTer domains), together with ßIV-spectrin SD (Figure 4G-J). The ankG SB domain colocalized with the submembrane ßIV-spectrin lattice on transverse sections (Figure 4G-H). The ankG CTer domain was found to be more intracellular than ßIV-spectrin: ankG CTer labeling was consistently found lining the intracellular side of the ßIV-spectrin lattice (Figure 4I). On radial profiles, this resulted in the ankG CTer intensity peaks being shifted toward the axoplasm compared to the ßIV-spectrin peaks (Figure 4J). As different AIS exhibit large variations in shapes (length, diameter, roundness), we devised a method to precisely quantify the radial arrangement of epitopes independently of these variations. A Gaussian curve was fitted on radial profiles, and the distance between the Gaussian maxima for the two channels was measured, providing the distance value relative to the ßIV-spectrin SD reference (Figure S5C-F). We determined the mean distance from the ßIV-spectrin SD to the extracellular epitope of neurofascin to be 17 ± 1 nm, and to the Nav intracellular loop to be 8 ± 1 nm (mean ± SEM, positive distance toward the axolemma, Figure 4M and Table S2). *α*-tubulin localization was broadly intracellular with a mean depth of −85 ± 4 nm (negative distance toward the axoplasm). The ankG SB domain was located just above ßIV-spectrin, with a mean distance of 6 ± 1 nm. The CTer domain of ankG was significantly more intracellular, with a mean distance of −26 ± 1 nm to ßIV-spectrin (Figure 4M). From these measurements, we determined ankG average radial extent: its CTer domain is on average 32 ± 1 nm deeper than its SB domain. We also mapped the position of the distal 480-kDa ankG tail relative to the ankG CTer (we could not use a ßIV-spectrin SD reference due to antibody host species issues). The two epitopes were colocalized on YZ sections (Figure 4K-L), showing that the distal tail localizes at the same depth as the CTer, with a relative distance close to zero (−2 ± 2 nm, Figure 4N). In conclusion, we have showed that the AIS is organized transversally, with the ankG carboxyterminus extending below the submembrane complex toward the axoplasm (Figure 4O).

### The radial extent of ankG is resistant to actin or microtubule perturbation

AnkG can be considered as a scaffold linker that binds both to actin via ßIV-spectrin, and to microtubules via EB1/3 proteins (Leterrier et al., 2011). Thus, we wondered if the ankG radial extent was dependent on the actin or microtubule cytoskeleton integrity. First, we used latA to acutely perturb actin, and measured the radial distance between the ankG CTer domain and the ßIV-spectrin SD. LatA treatment (5 μM, 1h) did not modify ankG arrangement: the ankG CTer domain was still found deeper than ßIV-spectrin (Figure 5A-D), with a distance of -26 ± 2 nm after treatment, compared to -24 ± 1 nm for vehicle (Figure 5E and Table S2). Thus, ankG radial extent does not depend on actin integrity.

**Figure 5.**
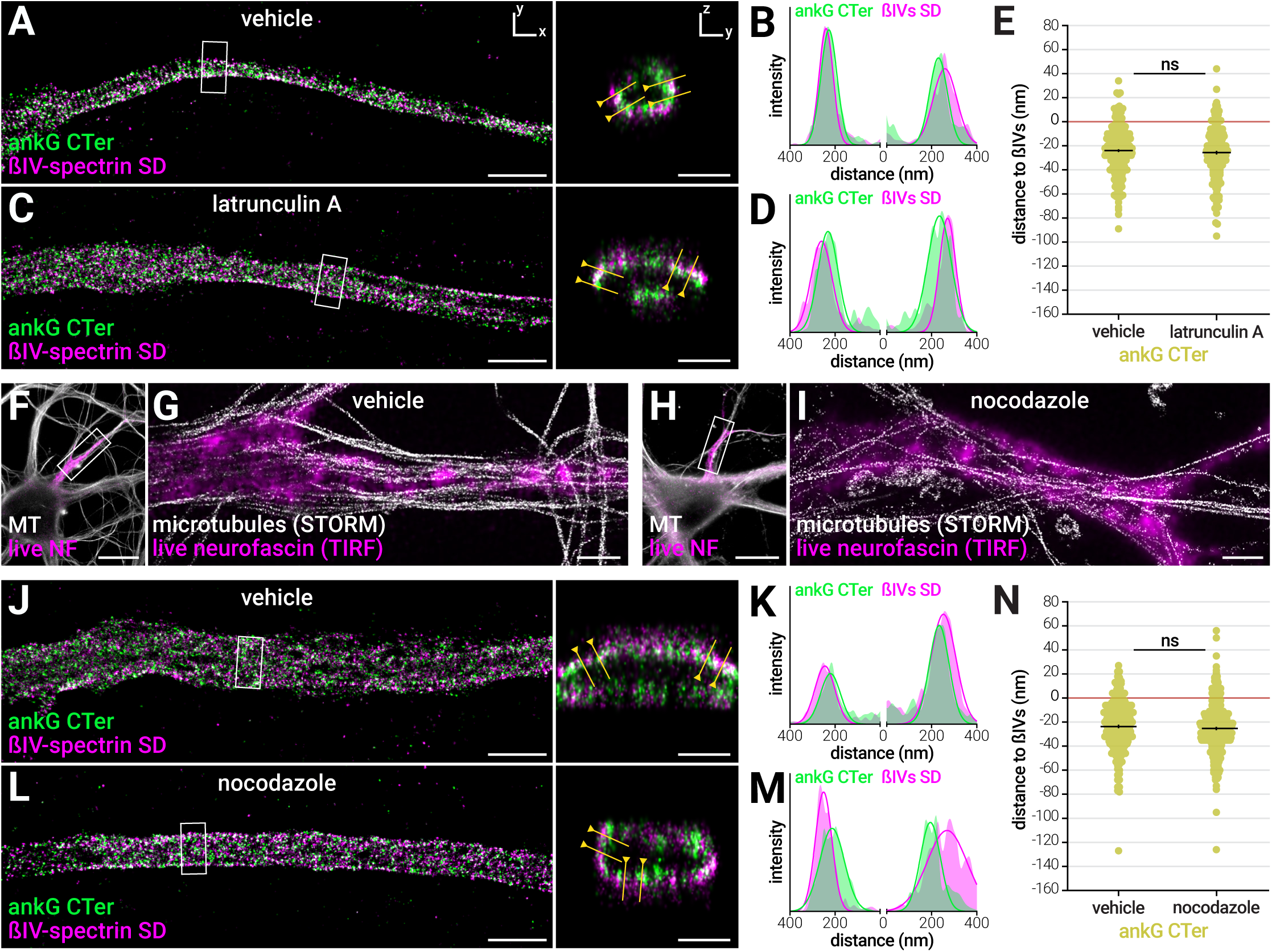
The radial extent of ankG is resistant to actin or microtubule perturbation (A) 3D-STORM image of the AIS from a neuron treated with vehicle (0.1% DMSO, 1h), fixed and labeled for ankG CTer (green) and ßIV-spectrin SD (magenta). Scale bars for (A, C, G, I, J, L) are 2 μm for XY image, 0.5 μm for YZ section. (B) Intensity profiles for each channel along the yellow line. (C-D) Same as (A-B), for an AIS from a sister culture treated with latrunculin A (latA, 5 μM, 1h). (E) Radial distance to ßIV-spectrin SD for the ankG CTer labeling in vehicle and latA-treated neurons (n=225-289 profiles, N=3). (F) Epifluorescence image of a neuron treated with vehicle (0.1% DMSO, 3h), labeled live for neurofascin (NF, magenta), fixed/extracted and labeled for microtubules (gray). (G) Overlay of a STORM images of the microtubule labeling (gray) with a scaled TIRF image of the live neurofascin labeling (magenta) along the AIS shown in (F). (H-I) Same as (F-G), for an AIS from a sister culture treated with nocodazole (20 μM, 3h). (J) 3D-STORM image of the AIS from a neuron treated with vehicle, fixed and labeled for ankG CTer (green) and ßIV-spectrin SD (magenta); (K) corresponding intensity profiles. (L-M) Same as (J-K), for an AIS from a sister culture treated with nocodazole. (N) Radial distance to ßIV-spectrin SD for the ankG CTer labeling in vehicle and nocodazole treated neurons (n=300-346, N=3-4). See also Figure S6.

Next, we used nocodazole at a high concentration (20 μM for 3h) to acutely perturb microtubules (Jaworski et al., 2009). At the diffraction-limited level, microtubules disassembly was observed after nocodazole treatment using an extraction/fixation procedure optimized for microtubule preservation (Figure S6A). A few filamentous structures were still brightly labeled, leading to a partial 55% decrease in labeling intensity at the AIS (13%, Figure S6B). However, STORM imaging showed that these remaining microtubules were running along, but not inside the AIS, likely belonging to distal axons (Figure 5F-I). In both vehicle and nocodazole treated neurons, periodicity measurement of the microtubule labeling along the AIS led to a flattened spacing histogram (190 ± 29 nm and 156 ± 32 nm, respectively) and low R squared values (0.18 ± 0.09 and 0.17 ± 0.11, respectively), providing a baseline for our measurement procedure (Figure S6C-J).

AIS components were barely affected by the nocodazole treatment at the diffraction level (12% drop in ankG concentration, no significant change for ßIV-spectrin or AIS length, Figure S6K-M). At the nanoscale level, the periodic ßIV-spectrin lattice was not perturbed by nocodazole treatment (Figure S6N-S), with no change in spacing spread for the ßIV-spectrin SD labeling (187 ± 8 nm for both vehicle and nocodazole, Figure S6P and S6S). This is in contrast with the partial disassembly of ßII-spectrin rings recently observed after nocodazole treatment (Zhong et al., 2014), confirming the specific robustness of the AIS scaffold. Finally, we mapped the radial organization of ankG after nocodazole treatment (Figure 5J-N). The localization of the ankG CTer domain was unaffected by nocodazole treatment: the ankG CTer domain distance to ßIV-spectrin was was -24 ± 1 nm for vehicle, and -25 ± 1 nm for nocodazole (Figure 5N). The radial arrangement of ankG, as well as the longitudinal periodicity of the ßIV-spectrin lattice, thus resists to actin filaments or microtubules perturbation.

### Elevated K+ partially disassemble the AIS, impairs the periodic lattice but not the ankG radial extent

To further probe the robustness of the AIS scaffold organization at the nanosTo further probe the robustness of the AIS scaffold organization at the nanoscale, we used a treatment that would affect the AIS morphology without completely disassembling it. Increase in intracellular calcium concentration has been implicated in the AIS morphological plasticity (Grubb and Burrone, 2010b) and shown to trigger AIS disassembly in ischemic injury (Schafer et al., 2009). We incubated neurons in the presence of 45 mM KCl for 3 hours (Redmond et al., 2002) to acutely increase intracellular calcium concentration. First, we measured the effect of these treatment on the overall morphology of the AIS at the diffraction-limited level (Figure 6A-B): high KCl treatment resulted in partial disassembly of the AIS, with a ~30% drop in ßIV-spectrin and ankG labeling intensity (Figure 6C) and a slight AIS shortening compared to the control condition (standard culture medium containing 5 mM KCl, Figure 6D). KCl treatment had a strong effect on the periodicity of the submembrane lattice (Fig 6E-H), with the ßIV-spectrin SD labeling exhibiting a flattened 189 ± 18 nm spacing distribution and a goodness of fit dropping from 0.58 ± 0.01 to 0.27 ± 0.01 (Figure 6I). In contrast, KCl treatments did not perturb the radial orientation of ankG (Figure 7J-N), with an average distance of ankG CTer to ßIV-spectrin SD at -24 ± 1 nm and -22 ± 2 nm for control and KCl treatments, respectively (Fig 7N). In conclusion, the range of perturbation performed (cytoskeleton, intracellular calcium) shows that the AIS nano-architecture is robust, with partial alterations appearing only when significant disassembly occurs for the whole AIS.

**Figure 6.**
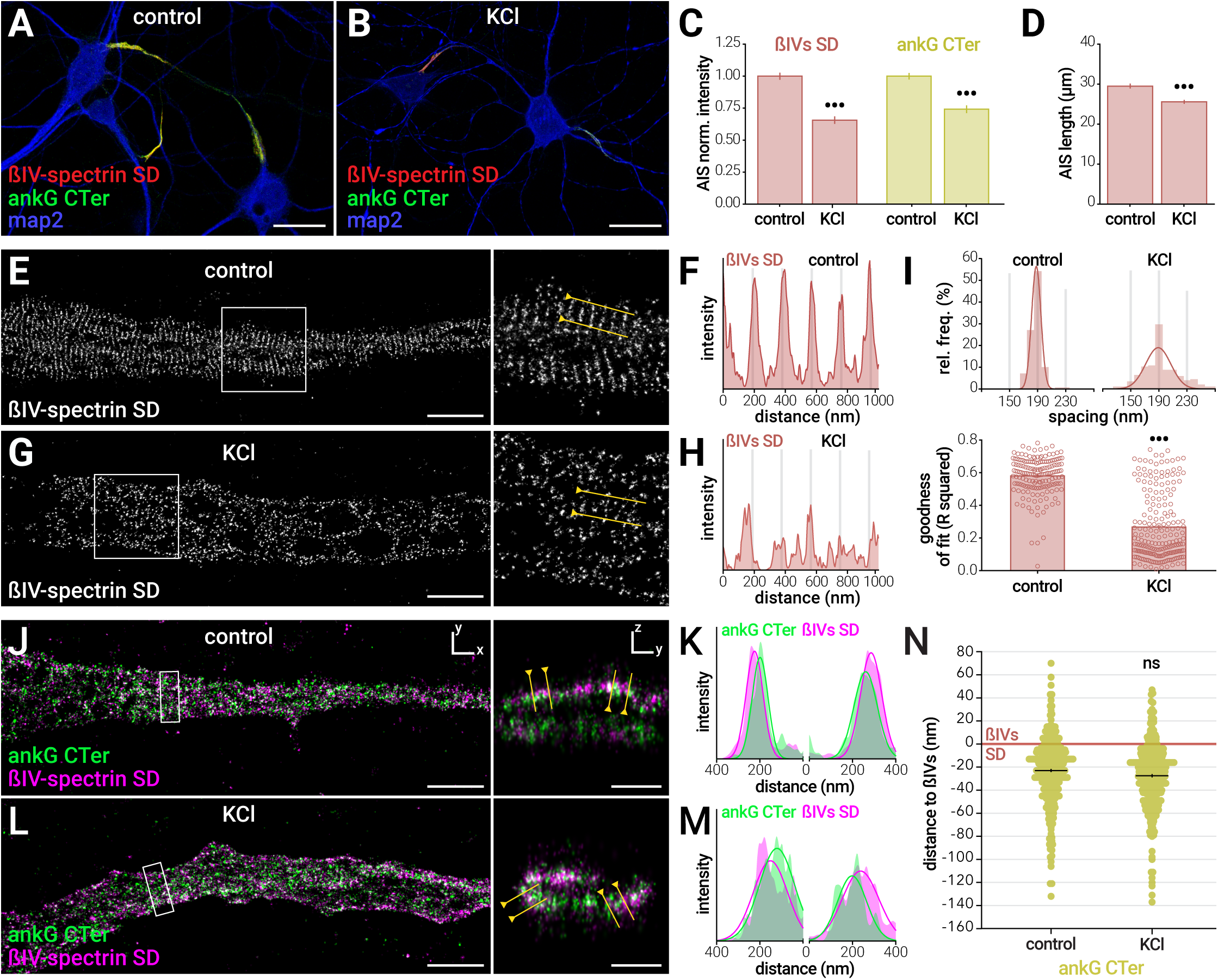
Elevated K+ partially disassemble the AIS, impairs the periodic lattice but not the ankG radial extent. (A-B) Deconvolved epifluorescence images of neurons untreated (A, control medium 5 mM KCI) or incubated with elevated K+ (B, 45 mM KCI, 3h), fixed and stained for ßIV-spectrin SD (red), ankG CTer (green) and map2 (blue). Scale bars, 20 μm. (C) Labeling intensity at the AIS of neurons for ßIV-spectrin SD (red, left) and ankG CTer (green, right), normalized to the vehicle condition. (D) AIS length measured on the ßIV-spectrin SD labeling (for C-D, n=103-126 AIS, N=3-4). (E) STORM image of AIS from control neurons labeled for ßIV-spectrin SD. Scale bars for (E, G, J, L) are 2 μm for XY images, 0.5 μm for YZ sections. (F) Intensity profile along the yellow line. (G-H) Same as (E-F) for an AIS from a sister culture treated with KCl (images and intensity profiles are identically processed for both conditions). (I) Top: histogram of the spacing values in the control and KCl conditions. Bottom: goodness of sinusoid fit (R squared) for the ßIV-spectrin SD labeling for control or KCl-treated neurons (for both graphs n=130-206 profiles, N=3-4). (J) 3D-STORM image of the AIS from control neurons, fixed and labeled for ankG CTer (green) and ßIV-spectrin SD (magenta). (K) Intensity profiles for each channel along the yellow lines. (L-M) Same as (J-K), for an AIS from a sister culture treated with KCl. (N) Radial distance to ßIV-spectrin SD for the ankG CTer labeling in control and KCl-treated neurons (n=415-431, N=3-4).

## Discussion

In this work, we have determined the nanoscale architecture of the AIS using super-resolution microscopy. STORM allowed us to quantitatively localize epitopes corresponding to known domains of AIS proteins, and infer the precise arrangement of the AIS scaffold. First, we directly demonstrate that the previously described ~190 nm periodic lattice along the AIS is formed by submembrane actin bands connected by longitudinal head-to-head ßIV-spectrin molecules. Furthermore, we reveal the specific resistance of this actin / ßIV-spectrin lattice against actin perturbation, compared to the ßII-spectrin-based lattice found along the distal axon. We characterized the yet unknown arrangement of the AIS master scaffolding protein ankG: the aminoterminal part of ankG associates with membrane proteins and ßIV-spectrin in the submembrane lattice, resulting in a periodic distribution. In contrast, the carboxyterminal part does not exhibit this periodicity: we explain this by demonstrating that the CTer of ankG localizes deeper in the axoplasm, below the submembrane lattice. Thanks to 3D-STORM, we measured a 32 nm radial extent between ankG SB and CTer domains. Finally, we showed that the lattice periodicity, as well as the radial extent of ankG, are resistant to a range of perturbations, revealing that this nanoscale organization is a robust and structural feature of the AIS.

### Longitudinal organization: periodic submembrane actin/ ßIV-spectrin lattice

The first striking feature of the AIS nano-architecture is the periodic arranThe first striking feature of the AIS nano-architecture is the periodic arrangement of its submembrane scaffold. We observed a regular distribution of actin and ßIV-spectrin extremities every ~190 nm along the AIS, confirming the periodic organization recently described (D’Este et al., 2015; Xu et al., 2013; Zhong et al., 2014). Purified brain spectrins have a length of ~ 195 nm (Bennett et al., 1982), suggesting a structural model where head-to-head ßIV-spectrin proteins connect to actin bands at their NT extremities, and to ankG/Nav channels complexes near their central SD domain (Figure 1U). Using 2-color STORM, we could detect correlated bands for actin / ßIV-spectrin NT, and anti-correlated ones for actin / ßIV-spectrin SD, directly demonstrating the validity of this model. Interestingly, two isoforms of ßIV-spectrin exist at the AIS: the 289-kDa ßIVΣ1, and the ~160 kDa ßIVΣ6 that lacks the aminoterminal part of ßIVΣ1 (Lacas-Gervais et al., 2004; Uemoto et al., 2007). ßIVΣ1 binds both actin and ankG and is likely to form the basis of the ~190 nm periodic lattice, in contrast to ßIVΣ6 that does not bind actin. The detection of the two isoforms by the SD antibody could explain the higher localization counts, and the larger individual bands compared to the NT antibody that only recognize ßIVΣ1.

Recent work showed that a periodic lattice of ßII-spectrin first appears along the whole axon, before being replaced at the AIS by ßIV-spectrin mature neurons used in our study (Zhong et al., 2014). Furthermore, we found the AIS ßIV-spectrin organization to be resistant to actin perturbation by latA (actin itself being partially stabilized at the AIS), as well as microtubule perturbation by nocodazole. It is likely that in the AIS, the ßIV-spectrin/actin complex is stabilized by the ankG/Nav channels complex. In the distal axon, the periodic ßII-spectrin lattice does not depend on the presence of its ankyrin B partner (Lorenzo et al., 2014), and this could explain why it is sensitive to cytoskeleton perturbation (Zhong et al., 2014). Ankyrin B presence is rather required for ßII-spectrin preferential concentration into the axon (Zhong et al., 2014), supporting a function in axonal transport rather than membrane structuration (Lorenzo et al., 2014).

### Transverse organization: radial extent of ankG

The second striking feature of the AIS nano-architecture is the radial arrangement of ankG. We demonstrate that the carboxyterminal part of ankG departs from the submembrane periodic lattice, and is found deeper in the axoplasm. 3D-STORM and quantification of the radial distribution of epitopes allowed us to measure the radial extent of ankG, with a 32 nm average distance between the SB and CTer domains (see Figure 4M). It is unlikely that antibody penetration was an issue in our mapping of ankG organization, as we could visualize microtubules at deeper positions, and sometimes throughout transverse sections of the AIS.

AnkG could stretch to hundreds of nanometers if its tail was fully extended and has been visualized as ~150 nm rods by scanning electron microscopy (Jones et al., 2014). The 32 nm radial extent value indicates that ankG actually lies in the vicinity of the plasma membrane, and may adopt convoluted conformations in the scaffold. This limited radial extent could also be adopted by the shorter 270-kDa ankG isoform that lacks the distal 1745 amino acids of the tail, as suggested by the similar depth measured using antibodies recognizing either the distal tail of 480-kDa ankG, or the CTer domain shared by both isoforms. Interestingly, the proximal part of the 480-kDa ankG specific tail (encompassing the S2417 residue) has recently been implicated in the recruitment of ßIV-spectrin to the AIS (P. M. Jenkins et al., 2015), suggesting it could be located in the submembrane lattice. We found that the distal part of the tail (antibody targeting residues 3516-3530) as well as the CTer domain were located ~25 deeper than ßIV-spectrin: this suggests that the ankG distal tail is the part that exits the submembrane lattice. The insertion of the large exon coding for the SR and tail domains of neuronal ankG has thus led to an extension of ankyrin G from the submembrane scaffold, potentially allowing the connection to more intracellular structures (Bennett and Lorenzo, 2013). Importantly, the intracellular localization of the ankG CTer domain was resistant to cytoskeleton perturbations. A short-term high KCl treatment that started to disassemble the AIS could alter the ßIV-spectrin periodic lattice, but not the radial extent of ankG. This nanoscale features of the AIS scaffold may thus be essential for the stability of the AIS as a compartment.

### Molecular organization of ankG hints at its role in regulating protein transport through the AIS

The “dendrification” of the proximal axon observed after ankG depletion (Hedstrom et al., 2008; Sobotzik et al., 2009) led to propose a role for the AIS in the maintenance of axonal identity via a diffusion barrier and an intracellular filter (Leterrier and Dargent, 2014; Rasband, 2010). Since its initial description (Song et al., 2009), the existence of an intracellular filter in the AIS that regulates protein traffic between the soma and axon has been a debated issue (Petersen et al., 2014; Watanabe et al., 2012). How ankG could participate in the regulation of protein transport into the axon remains unknown, as ankG has been primarily detected near the plasma membrane by immunogold electron microscopy (Iwakura et al., 2012; Le Bras et al., 2013). Even if ankG forms ~150 nm long rods (Jones et al., 2014), our results show that its reaches depths within ~50 nm of the plasma membrane, ruling out a fully radial orientation and the hypothesis of ankG extending through the axoplasm to reach deep intracellular targets (Davis et al., 1996; Leterrier and Dargent, 2014). The shallow depth reached by ankG in our study suggest that the AIS scaffold could rather recruit and spatially organize a population of microtubules close to the axolemma (Westrum and Gray, 1976). This specific microtubule organization could in turn influence polarized traffic to and from the axon, explaining how the AIS participate in the sorting of vesicular trafficking. Interestingly, a recently identified giant ankG isoform organizes the distribution of microtubules along the axon in Drosophila neurons (Stephan et al., 2015).

### The AIS, from cellular traffic to brain and nervous system disorders

In conclusion, we show that the AIS scaffold is a precisely organized compartment, with a longitudinal periodicity and a radial layering. This nanoscale organization is robust, pointing to its potential importance for proper cell function. Our work confirms the strength of super-resolution microscopy for elucidating the architecture of neuronal assemblies, down to the macromolecular level (Maglione and Sigrist, 2013). Knowledge of the AIS nanoscale architecture will help deciphering the molecular mechanisms that underpin neuronal excitability and protein mobility in polarized cells. Beyond this fundamental relevance, understanding the AIS structure and function has neuro-pathological implications. AnkG gene variants and mutations have been consistently associated with in several neuropsychiatric disorders, including bipolar disorders and schizophrenia (Iqbal et al., 2013; Leussis et al., 2012). Furthermore, affected axonal transport is a key factor in most neurodegenerative diseases, where the gatekeeper function of the AIS was recently implicated (Sun et al., 2014). Our structural work will hopefully open the way to a better understanding of the AIS crucial functions, in physiological as well as pathological contexts.

## Materials and Methods

### Antibodies, plasmids and reagents

Rabbit polyclonal anti ßIV-spectrin antibodies were gifts from Matthew Rasband (Baylor College of Medicine, Austin, TX). Rabbit polyclonal anti 480-kDa ankG (residues 3516-3530 of human 480-kDa ankG) was a gift from François Couraud (Université Pierre et Marie Curie, Paris). For two-color STORM, paired fluorophore-conjugated secondary antibodies were made by coupling unconjugated antibodies with reactive activator and reporter fluorophores, according to the N-STORM sample preparation protocol (Nikon Instruments).

### Animals and neuronal cultures

The use of Wistar rats followed the guidelines established by the European Animal Care and Use Committee (86/609/CEE) and was approved by the local ethics committee (agreement D13-055-8). Rat hippocampal neurons were cultured on 18 mm coverslips at a density of 6,000/cm2 following the Banker method, above a feeder glia layer in B27-supplemented medium (Kaech and Banker, 2006).

### Immunocytochemistry and STORM imaging

After 14 to 21 days in culture, neurons were fixed using 4% PFA or using an extraction-fixation method optimized for microtubule labeling. After blocking, they were incubated with primary antibodies overnight at 4°C, then with secondary antibodies for 1h at room temperature. STORM imaging was performed on an N-STORM microscope (Nikon Instruments). Coverslips were imaged in STORM buffer: Tris 50 mM pH 8, NaCl 10 mM, 10% glucose, 100 mM MEA, 3.5 U/mL pyranose oxidase, 40 μg/mL catalase. For single color imaging, the sample was continuously illuminated at 647 nm (full power) and a 30,000-60,000 images were acquired at 67 Hz, with progressive reactivation by simultaneous 405 nm illumination. Two-color STORM imaging was either performed by successive imaging with 488 and 647 nm lasers (for double staining comprising phalloidin-Atto 488), or using alternated sequences of one activator frame followed by three reporter frames. The N-STORM software (Nikon Instruments) was used for the localization of single fluorophores.

### Image Processing and analysis

Image reconstructions were performed using the ThunderSTORM ImageJ plugin (Ovesny et al., 2014). Custom scripts and macros were used to automate images reconstructions. For quantification of longitudinal periodicity, intensity profiles were fitted using a sinusoid function. The histogram of spacing values was fitted with a Gaussian curve to obtain the mean spacing and spread. For quantification of radial distributions, radial profiles were obtained on YZ transverse projections, and the distance between epitopes defined as the difference between the maxima of Gaussian fits on each channel. For quantification of the AIS length and components intensity on epifluorescence images, a threshold was applied on line profiles along the axon to detect the AIS position (Grubb and Burrone, 2010b). Significances were tested using two-tailed unpaired t-tests (two conditions) or one-way ANOVA followed by Tukey post-test (3 or more conditions). In all Figures and tables significance is coded as: ns non-significant,• p < 0.05, •• p < 0.01, ••• p < 0.001.

## Acknowledgments

We thank M. Rasband, V. Bennett and F. Couraud for providing antibodies and plasmids. We also would like to thank D. Choquet and J.-B. Sibarita for insightful discussions; D. Marguet, S. Mailfert and M. Mondin for their help during the initial stages of this study; S. Bezin at Nikon Instruments and M.-P. Blanchard at CRN2M imaging facility for their assistance; S. Roy, F. Castets and M.-J. Papandreou for discussions and careful reading of the manuscript.

## Author contributions

CL: conception and design, acquisition of data, analysis and interpretation of data, drafting or revising the article. JP: acquisition of data, analysis and interpretation of data. GC: acquisition of data. CD: acquisition of data. FRB: contributed essential data or reagents. BD: conception and design, interpretation of data, drafting or revising the article. This work was supported by a grant to BD from the French Agence Nationale de la Recherche (ANR-2011-BSV4-001-1). The authors declare that no competing interests exist.

## Supplementary Figures

**Figure S1.**
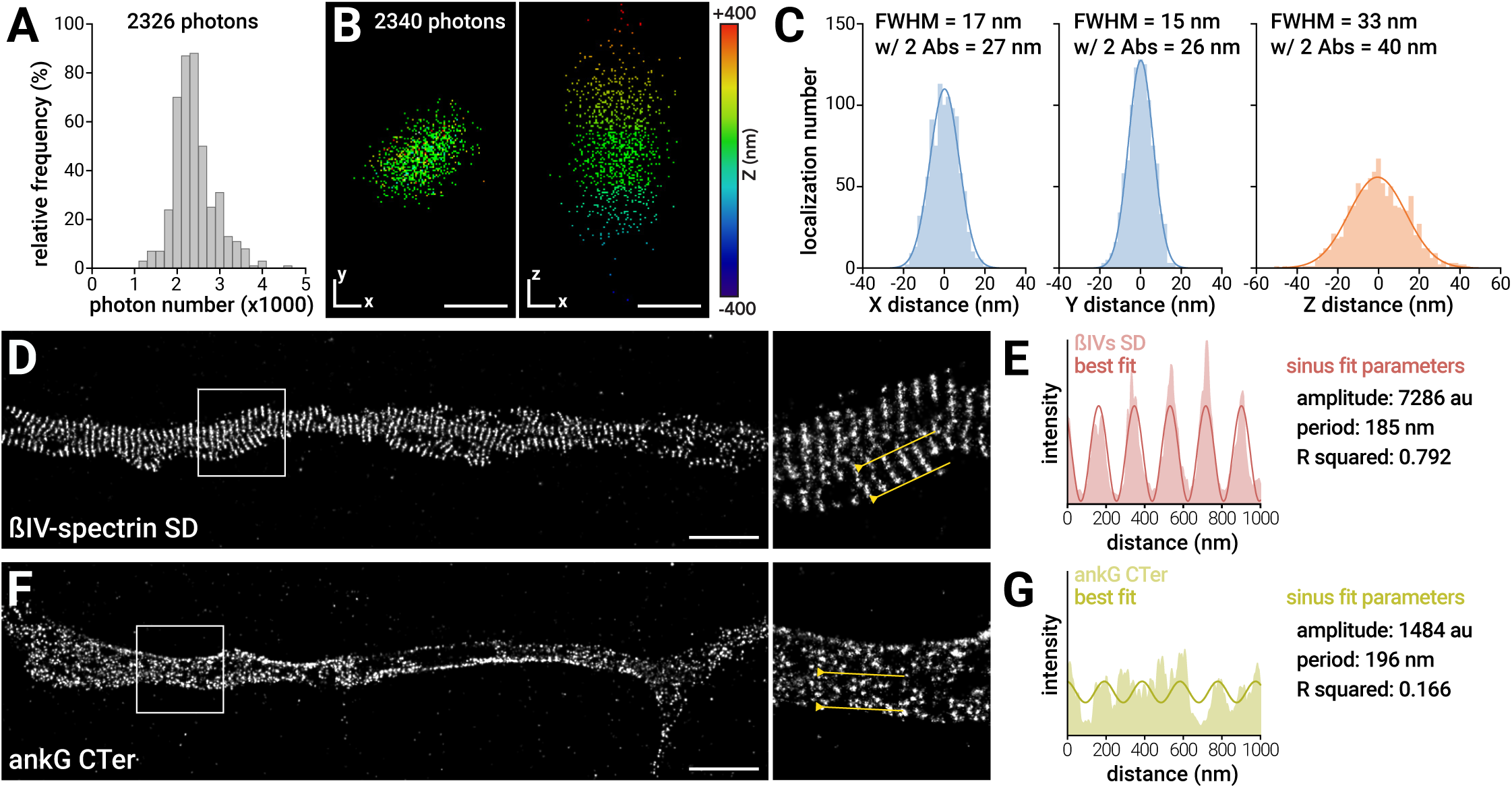
Localization precision of STORM images and quantification procedure for periodicity measurements, related to Figure 1. (A-C) Measurement of the STORM images localization precision. (A) Histogram of the median number of photons emitted by localizations from each single image from n=429 STORM images used in this work. Average number is 2326 photons. (B) 1000 frames of a 0.1 μm fluorescent bead were acquired, with laser power adjusted so that median number of photons emitted on each frame was 2340. The resulting localization scatter from the localization software is plotted as XY (left) and XY (right) projections, color coded for depth. Scale bars, 20 nm. (C) Gaussian fits of the localizations scatter provide an accurate measure of the localization precision, expressed as full width at half maximum (FWHM) above each X, Y and Z curve. The second “w/ 2 Abs” value takes into account the added uncertainty from two 15 nm-sized antibodies (see Extended Experimental Procedures). (D-G) Quantification of the labeling periodicity on STORM images. (D) STORM image of an AIS with a highly periodic labeling (ßIV-spectrin SD). Scale bars for (D, F) are 2 μm. (E) Intensity profile along the yellow line (light red); red curve is the sinusoid fitted on the profile, with fitted parameters given on the right. (F-G) Same as D-E, with a low periodicity labeling (ankG CTer domain, green on profile).

**Figure S2.**
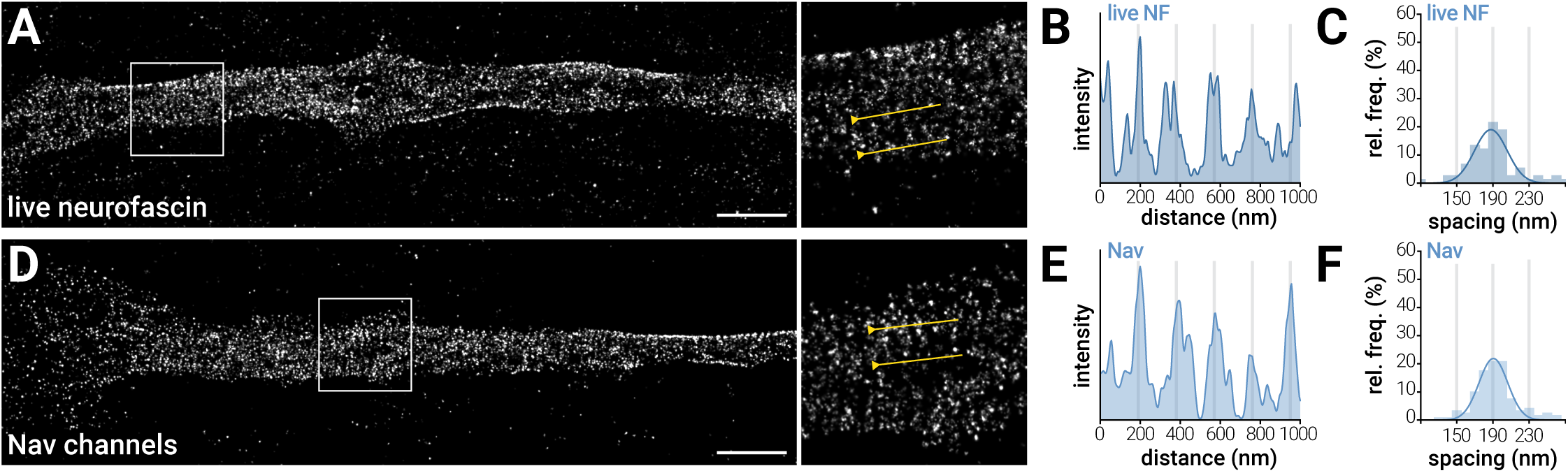
Detection of a periodic arrangement of neurofascin and Nav channels at the AIS, related to Figure 1. (A) STORM image of an AIS labeled live for neurofascin. Scale bars for (A, D) are 2 μm. (B) Intensity profile along the yellow line. (C) Histogram of the spacing values (n=74 profiles, N=3). (D) STORM image of an AIS labeled for sodium channels; (E) corresponding intensity profile; (F) histogram of spacing values (n=120, N=3).

**Figure S3.**
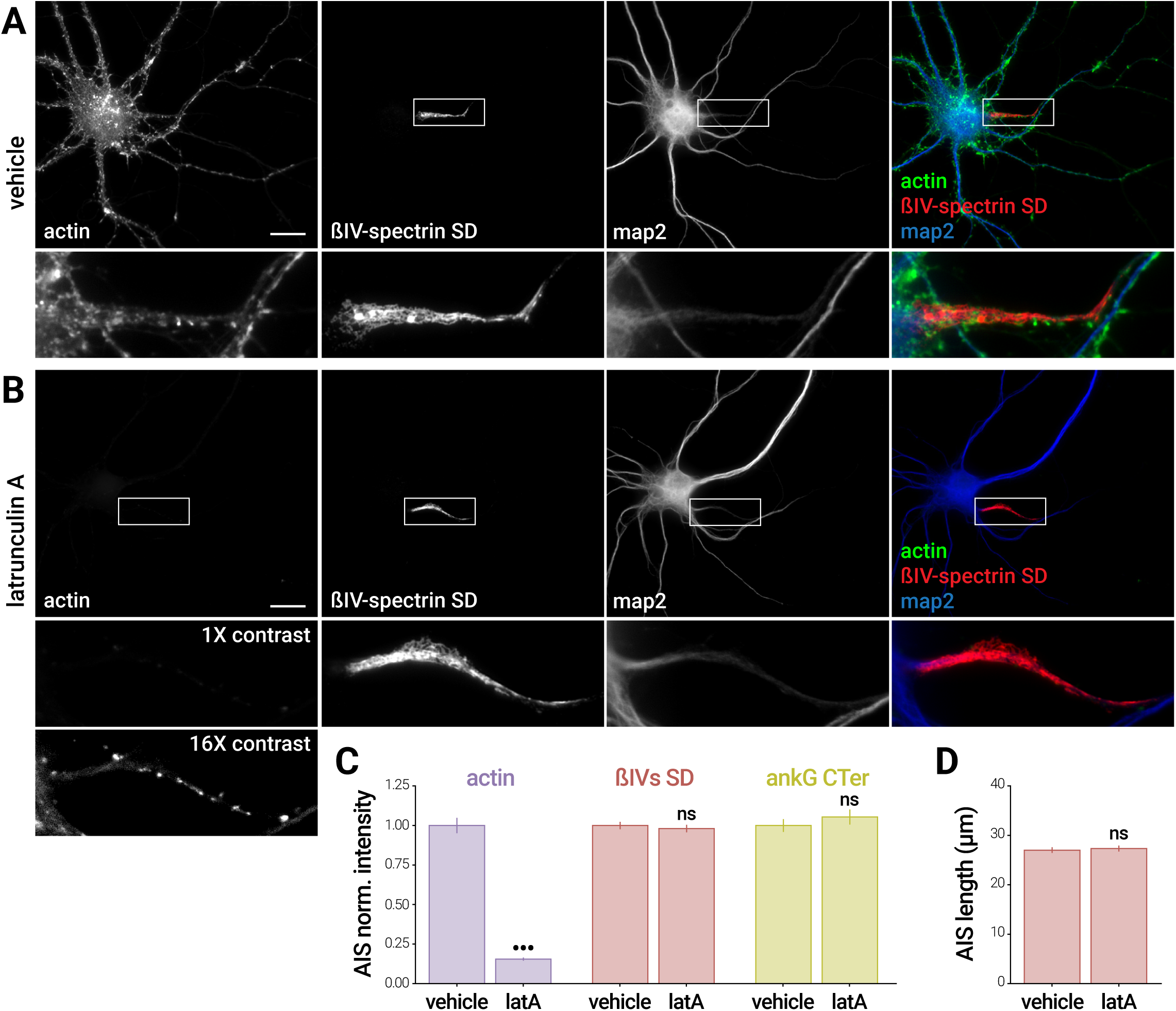
Effect of latrunculin A treatment on actin and AIS components, related to Figure 2. (A-B) Widefield epifluorescence images of neurons treated with vehicle (A, DMSO 0.1%, 1h) or latA (B, 5 μM, 1h), fixed and labeled for actin (phalloidin, green on overlay), ßIV-spectrin SD (red on overlay) and map2 (blue on overlay). For the actin single channel image of the latA-treated neuron, the zoom is shown with the same contrast as the vehicle-treated neuron (top box) or with a 16X enhanced contrast to reveal residual actin (bottom box). Scale bars, 20 μm. (C) Labeling intensity at the AIS of treated neurons for actin (purple), ßIV-spectrin SD (red) and ankG CTer (green), normalized to the vehicle condition. (D) AIS length measured on the ßIV-spectrin SD labeling (for C, D n=68-159 AIS, N=2-4).

**Figure S4.**
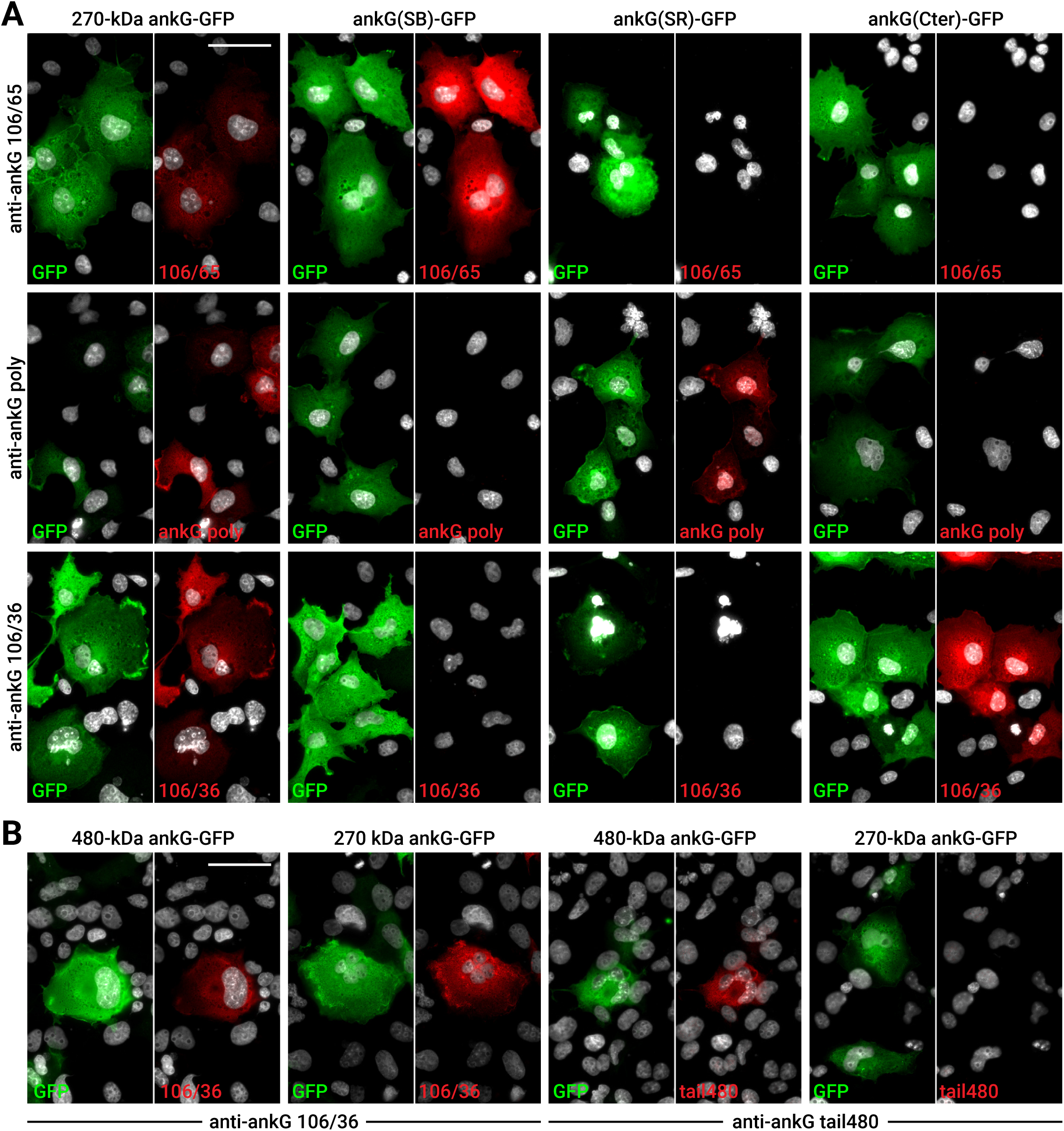
Determination of the ankG domains targeted by the different anti-ankG antibodies, related to Figure 3. COS cells expressing 480-kDa ankG-GFP, 270-kDa ankG-GFP or different isolated ankG domains fused to GFP (green) were labeled using one of the four anti-ankG antibodies used in this study (red). (A) Among the three antibodies that recognize the both 480-kDa and 270-kDa ankG-GFP (left column), the 106/65 antibody recognizes the SB domain, the polyclonal antibody recognizes the SR domain, and the 106/36 antibody recognizes the CTer domain. (B) Both 480- and 270-kDa ankG-GFP are recognized by the 106/36 antibody targeting the CTer domain (left). Only the 480-kDa ankG-GFP, not the 270-kDa ankG-GFP, is recognized by the anti-ankG tail480 (right). Scale bars, 50 μm.

**Figure S5.**
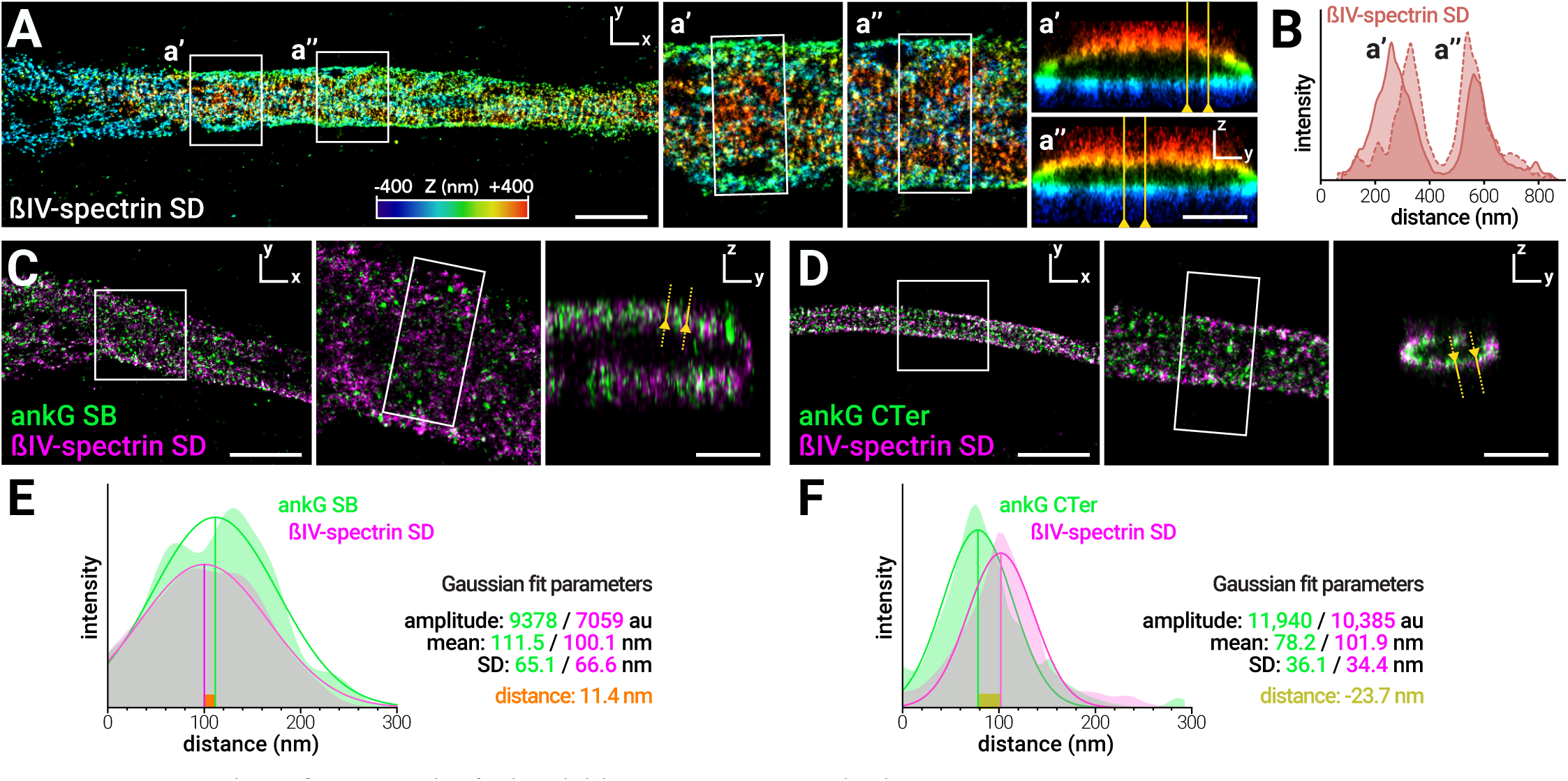
3D-STORM and quantification procedure for the radial distance measurements, related to Figure 4. (A) Image of an AIS labeled for ßIV-spectrin SD (color-coded for depth) demonstrating 3D-STORM imaging. Scale bar is 2 μm for XY image, 0.5 μm for YZ sections. (B) Intensity profile along the yellow lines on the transverse sections (A, a’ and a”). (C-F) Quantification of radial distribution on 3D-STORM images. (C-D) STORM images of an AIS labeled for ankG SB (C, green) and ßIV-spectrin SD (magenta) or ankG CTer (D, green) and ßIV-spectrin SD (magenta). Scale bar is 2 μm on XY image, 0.5 μm on YZ section. (E-F) Examples of quantification of the radial distribution for labeling shown in (C-D). Filled curves are intensity profiles for each channel along the yellow line on YZ sections in (C) and (D). Curves are Gaussian fit of the intensity profiles: ankG SB (green) and ßIV-spectrin SD (magenta) for (E), ankG CTer (green) and ßIV-spectrin SD (magenta) for (F). From the fit parameters (given on the right), the radial distance is calculated by subtracting the means of the Gaussian fits for the two channels: positive radial distance (E) indicates that the ankG SB labeling localizes at the periphery of the ßIV-spectrin SD labeling, whereas negative radial distance (F) indicates that the ankG CTer labeling is found inside the ßIV-spectrin SD labeling.

**Figure S6.**
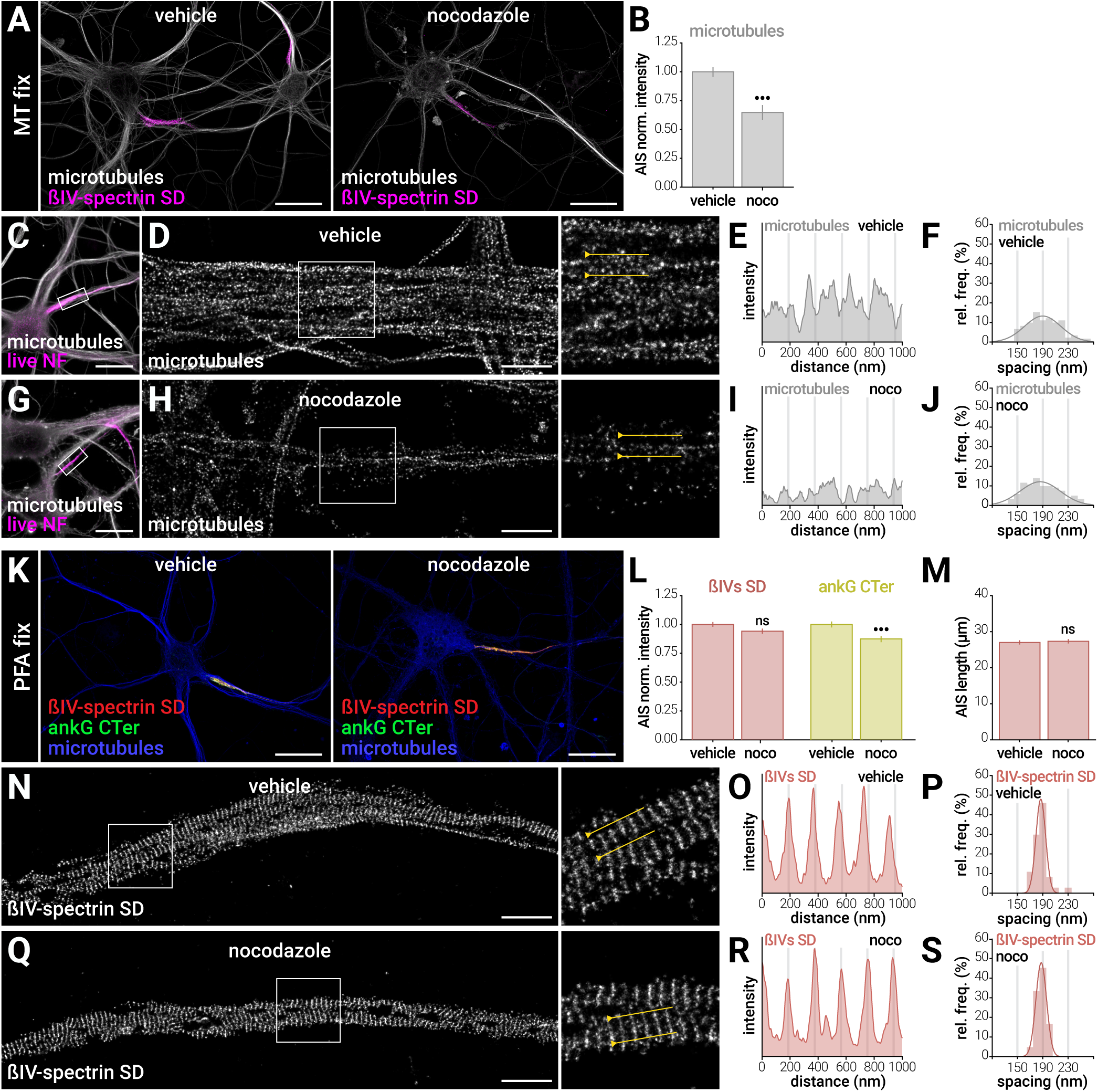
Effect of nocodazole treatment on microtubules and AIS components, related to Figure 5. (A) Deconvolved epifluorescence image of neurons treated with vehicle (left, DMSO 0.1%, 3h) or nocodazole (right, 20 μM, 3h), extracted/fixed (MT fix) and stained for micro-tubules (gray) and ßIV-spectrin SD (magenta). Scale bars for (A, C, G, K) are 20 μm. (B) Labeling intensity for the microtubule labeling at the AIS of treated neurons, normalized to the vehicle condition (for B, L, M, n=111-116 AIS, N=3). (C) Epifluorescence image of a neuron treated with vehicle, labeled live for neurofascin (NF, magenta), fixed/extracted and labeled for microtubules (gray). (D) STORM image of the microtubule labeling along its AIS; (E) intensity profile along the yellow line on the zoomed STORM image; (F) histogram of spacing values (n=183 profiles N=4). Scale bars for (D, H, N, Q) are 2 μm. (G-J) Same as (C-F), for a neuron from a sister culture treated with nocodazole (histogram, n=122 N=4). Intensity profile (I) processed identically to the vehicle condition (E). (K) Deconvolved epifluorescence image of neurons treated with vehicle (left) or nocodazole (right), fixed (PFA fix) and stained for ßIV-spectrin SD (red), ankG CTer (green) and microtubules (blue). (L) Labeling intensity at the AIS of treated neurons for ßIV-spectrin SD (red, left) and ankG CTer (green, right), normalized to the vehicle condition. (M) AIS length measured on the ßIV-spectrin SD labeling.

**Table S1.**
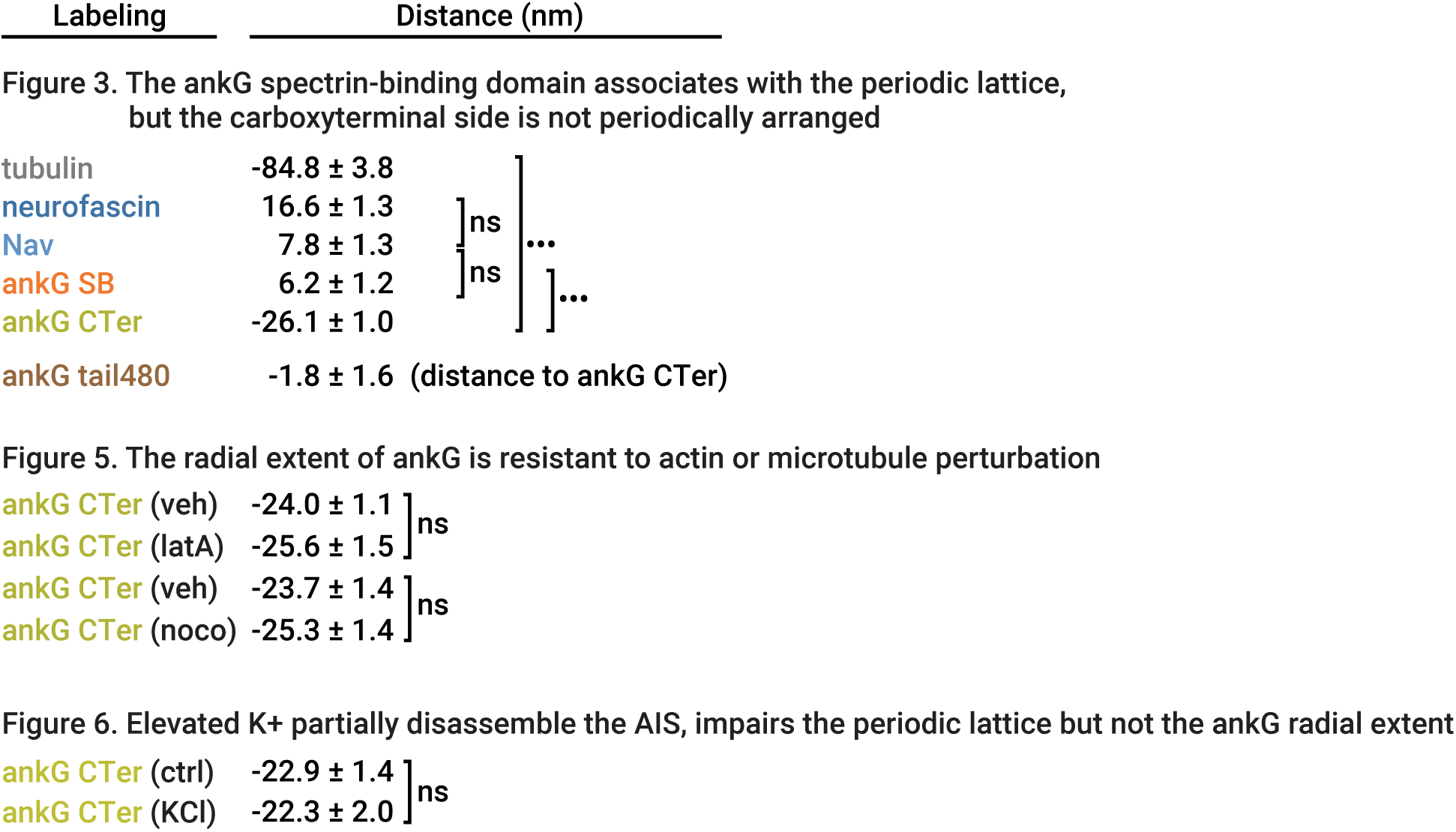
Spacing and periodicity measurements. The distribution of spacing values obtained by fitting sinusoids on intensity profiles is fitted by a Gaussian curve. The mean of this Gaussian is reported in the “Mean spacing” column, and the standard deviation of this Gaussian is reported in the “Spread” column. Errors are obtained from the fit procedure. The “R squared” column shows values for the goodness of sinusoid fit for all profiles in a given condition. All values are mean ± SEM. * same dataset (actin treated with vehicle) used in both figures.

**Table S2.**
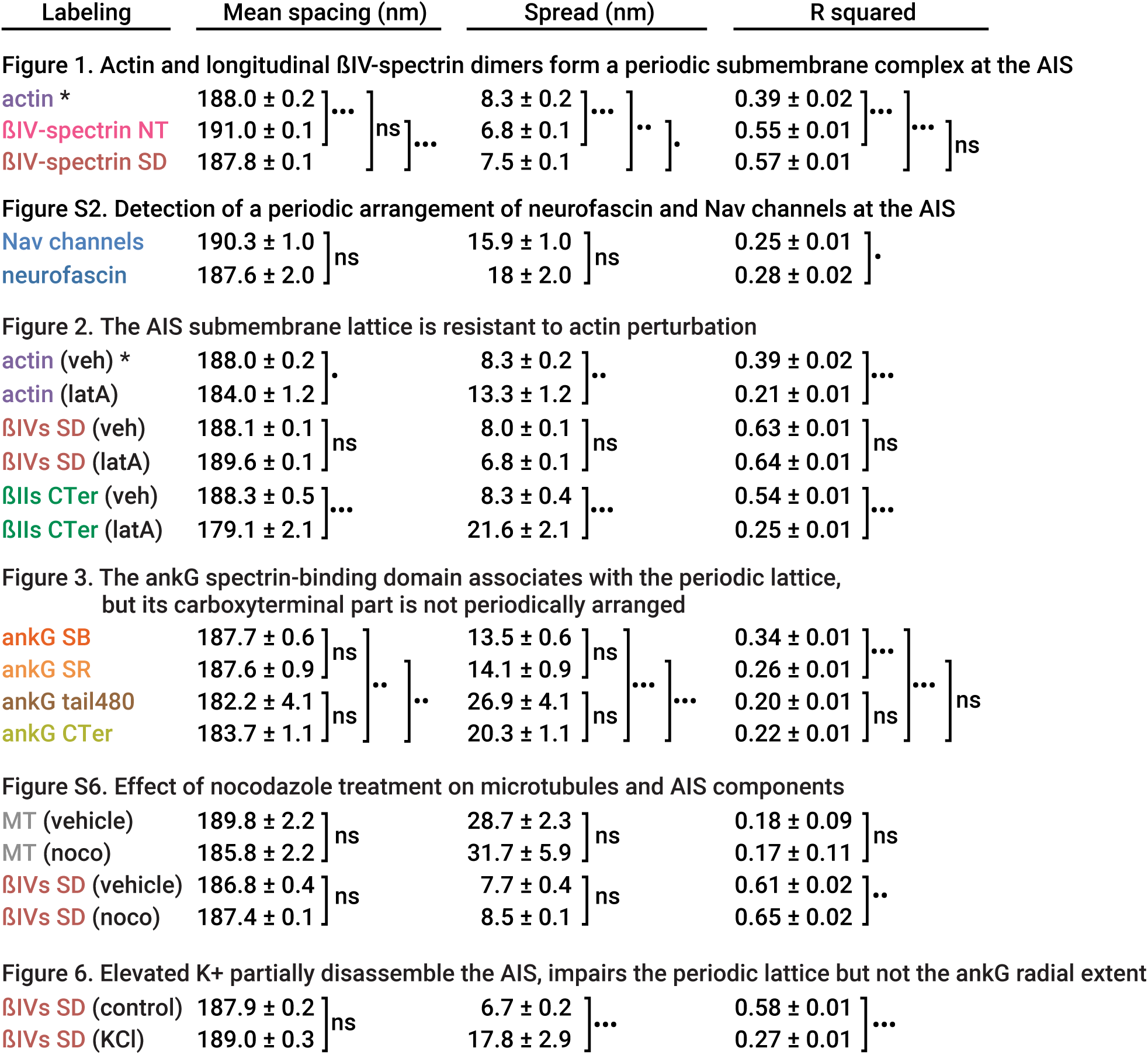
Radial distance measurements. These distances are mean ± SEM obtained from measurements and fit on radial intensity profiles. Distance is to ßIV-spectrin SD labeling, unless specified otherwise.

## Extended Experimental Procedures

### Antibodies, plasmids and reagents

Rabbit polyclonal anti ßIV-spectrin antibodies (against residues 15-38 and 2237-2256 of human ßIV-spectrin Σ1 for the NT and SD antibodies, respectively) were gifts from Matthew Rasband (Baylor College of Medicine, Austin, TX). Mouse monoclonal anti pan-Nav channels (against the intracellular III-IV loop, clone K58/36, #S8809) and mouse monoclonal anti *α*-tubulin (clone DM1a, #T3199) were from Sigma. Rabbit polyclonal anti ankG was generated against residues 1633-1650 of human ankG (SR domain)(Bréchet et al., 2008). Mouse monoclonal anti ßII-spectrin (against residues 2101-2189 of human ßII-spectrin) was from BD Biosciences (#612563). Mouse monoclonal antibodies anti ankG (against rat SB and CTer domains, clone 106/65 and 106/36, respectively), and anti neurofascin (against extracellular domain residues 25-1110 common to rat 155- and 186-kDa neurofascin, clone A12/18) were from NeuroMab. Rabbit polyclonal anti 480-kDa ankG (residues 3516-3530 of human 480-kDa ankG) was a gift from François Couraud (Université Pierre et Marie Curie, Paris). Rabbit polyclonal anti *α*-tubulin was from abcam (#18251).

Donkey and goat anti-rabbit and anti-mouse secondary antibodies conjugated to Alexa Fluor 488 and 555 were from Life Technologies, secondary antibodies conjugated to DyLight 405 and Alexa Fluor 647 from Jackson ImmunoResearch. Paired fluorophore-conjugated donkey anti-mouse and anti-rabbit secondary antibodies were made by coupling unconjugated antibodies (Jackson ImmunoResearch) with Alexa Fluor 405 carboxylic acid, succinimidyl ester (#A30000, Life Technologies) or Cy3 mono-reactive dye (#PA23001, GE Healthcare) as activator and Alexa Fluor 647 carboxylic acid succinimidyl ester (#A20006, Life Technologies) as reporter, according to the N-STORM sample preparation protocol (Nikon Instruments).

480-kDa ankG-GFP, 270-kDa ankG-GFP and individual ankG domains were gifts from Vann Bennett (Duke University, Durham, NC)(Jenkins et al., 2015; Zhang and Bennett, 1998). Alexa Fluor 647-conjugated phalloidin was from Life Technologies (#A22287); Atto 488-conjugated phalloidin (#49409), latrunculin A (#L5163, stock 5mM in DMSO), nocodazole (#M1404, stock 20 mM in dimethyl sulfoxide [DMSO]), glutaraldehyde (#G5882, 25% in water), ß-mercaptoethylamine ([MEA], #30070, stock 1 mM in HCl 360 mM), pyranose oxidase (#P4434, 200U/mL in GOD buffer/glycerol) and catalase (#C40, 5 mg/mL in GOD/glycerol) were from Sigma. GOD buffer is 24 mM PIPES, 4 mM MgCL2, 2 mM EGTA, pH 6.8. Paraformaldehyde ([PFA], #15714, 32% in water) was from Electron Microscopy Sciences.

### Animals and neuronal cultures

The use of Wistar rats followed the guidelines established by the European Animal Care and Use Committee (86/609/CEE) and was approved by the local ethics committee (agreement D13-055-8). Rat hippocampal neurons were cultured on 18 mm, #1.5H coverslips at a density of 6,000/ cm2 following the Banker method, above a feeder glia layer in B27-supplemented medium (Kaech and Banker, 2006).

### Immunocytochemistry

After 14 to 21 days in culture, neurons were fixed using 4% PFA in 0.1 M phosphate buffer saline (PB) for 10 minutes at room temperature (RT). For labeling of microtubules (Figure S6), neurons were fixed and extracted with 0.25% glutaraldehyde, 0.3% Triton X-100 in extraction buffer (PIPES 80 mM pH 6.9, 150 mM NaCl, 4 mM MgCl2, 1 mM EGTA, 5 mM glucose) for 1 minute followed by 4% PFA, 4% sucrose in PB for 10 minutes (Yau et al., 2014). Anti ßIV-spectrin SD labeling was partially retained after this extraction/fixation procedure, allowing to identify the AIS. Alternatively, neurons were labeled live before extraction and fixation using an anti-neurofascin antibody recognizing an extracellular epitope of 186-KDa neurofascin for 7 minutes at 37°C.

After rinses in PB, neurons were blocked for 1h at RT in immunocytochemistry buffer (ICC: 0.22% gelatin, 0.1% Triton X-100 in PB), and incubated with primary antibodies diluted in ICC overnight at 4°C. After rinses in ICC, neurons were incubated with secondary antibodies diluted in ICC for 1h at RT, and finally rinsed and kept in PB + 0.02% sodium azide at 4°C before STORM imaging. For STORM imaging of actin, phalloidin was added at 0.5 μM in PB for 1h just before proceeding to imaging (Xu et al., 2013). For epifluorescence imaging, coverslips were mounted in ProLong Gold (Life Technologies).

### Epifluorescence microscopy

Diffraction-limited images were obtained using an Axio-Observer upright microscope (Zeiss) equipped with a 40X NA 1.4 or 63X NA 1.46 objective and an Orca-Flash4.0 camera (Hamamatsu) and. Appropriate hard-coated filters and dichroic mirrors were used for each fluorophore. Quantifications were performed on single, unprocessed 40X images. An Apotome optical sectioning module (Zeiss) and post-acquisition deconvolution (Zen software, Zeiss) were used to acquire and process images used for illustration.

### STORM imaging

STORM imaging was performed on an N-STORM microscope (Nikon Instruments). Coverslips were mounted in a Ludin Chamber (Life Imaging Services) and imaged in STORM buffer: Tris 50 mM pH 8, NaCl 10 mM, 10% glucose, 100 mM MEA, 3.5 U/mL pyranose oxidase, 40 μg/mL catalase. The N-STORM system uses an Agilent MLC-400B laser launch with 405 nm (50 mW maximum fiber output power), 488 nm (80 mW), 561 mW (80 mW) and 647 nm (125 mW) solid-state lasers, a 100X NA 1.49 objective and an Ixon DU-897 camera (Andor). After locating a suitable neuron using low-intensity illumination, a TIRF image was acquired, followed by a STORM acquisition. For single color imaging, the sample (stained with an Alexa Fluor 647 conjugated secondary antibody) was continuously illuminated at 647 nm (full power) and a series of 30,000-60,000 images (256x256 pixels, 15 ms exposure time) was acquired. Reactivation of the fluorophore was performed during acquisition by increasing illumination with the 405 nm laser.

For direct two-color STORM imaging (Figure 1K-P), Alexa 647-conjugated antibody labeling was imaged first in STORM buffer using 647 nm laser illumination. Medium was then exchanged with a saline buffer (Tris 50 mM pH 8, NaCl 10 mM) for imaging the Atto 488-conjugated phalloidin labeling using 488 nm laser excitation (Nanguneri et al., 2014), and the two channels were aligned using fiducial beads (Tetraspeck 0.1 μm, Life Technologies #T7279). For two-color imaging using secondary antibodies labeled with activator-reporters fluorophore pairs (Alexa Fluor 405-Alexa Fluor 647 and Cy3-Alexa Fluor 647), the sample was illuminated using sequences of one activator frame (405 or 561 nm) followed by three reporter frames (647 nm) (Bates et al., 2007). For 3D-STORM, a cylindrical lens was introduced in the optical path, leading to a Z-dependent asymmetry of the PSF (Huang et al., 2008). The N-STORM software (Nikon Instruments) was used for the localization of single fluorophore activations. After filtering only localizations with more than 900 photons, the list of localizations was exported as a text file.

### Spatial precision of the STORM imaging

To evaluate the localization precision of STORM in our hands, we used a method similar to Nair et al. (Nair et al., 2013) (Figure S1A-C). As the localization precision directly depends on the number of photons emitted during a single fluorophore activation, we measured the median number of photons for all localizations from 429 STORM images and obtained an average of 2320 photons. We continuously imaged 100 nm fluorescent beads at an excitation power (2% 647 nm laser) resulting in a similar number of photons (2340) emitted per frame for 1000 frames. Beads were localized in 3D using the N-STORM software. The spread of the beads localizations on successive frames directly gives the localization precision: we obtained a SD of 7, 6 and 14 nm corresponding to full width at half maximum [FWHM] of 17, 14 and 33 nm in X, Y and Z, respectively (Figure S1A).

Beyond the fluorophore localization precision, the use of a combination of primary and secondary antibodies to detect endogenous epitopes can degrade the precision of epitope localization, as discussed previously (Dani et al., 2010). Each antibody is ~15 nm in size, adding ~6.5 nm of additional uncertainty (SD of a Gaussian with a 15 nm FWHM). Primary and secondary antibody layers will add to the localization SD measured above as √(7ˆ2 + 6.5ˆ2 + 6.5ˆ2) = 11.0 nm in X, 11.6 nm in Y and 16.7 nm in Z. This corresponds to FWHM of 26, 27 and 39 nm in X, Y and Z, respectively. Finally, the high number of profiles n used for quantifications (several hundreds) allows to narrow the error on the average position of epitopes by a 1/√n factor, bringing it to a few nanometers. Accordingly, even the small radial distances such as the one between ankG SB domain and ßIV-spectrin CTer (<10 nm, see Figure 4) were consistently reproduced in each independent experiment.

### Image Processing and analysis

Image reconstructions were performed using the ThunderSTORM ImageJ plugin (Ovesny et al., 2014) in Fiji software (Schindelin et al., 2012). Custom scripts and macros were used to translate localization files from N-STORM to ThunderSTORM formats, as well as automate images reconstruction for whole images, detailed zooms and YZ transverse projections.

Quantification of the longitudinal periodicity (see Figure S1D-G and Table S1) was performed on 2D or projected 3D-STORM images using a custom ImageJ script. 3 to 5 regions (3x3 μm) were selected along an AIS, and high resolution reconstructions (4 nm/pixel) were generated. Two intensity profiles (200 nm wide for ßII-spectrin, 400 nm for others) were drawn on each reconstruction where periodicity was best seen by eye, and fitted using a sinusoid function with the spacing defined as the period P, seeded at 185 nm. The histogram of spacing values was fitted with a Gaussian curve (in Prism software): mean spacing correspond to the Gaussian mean, and spread to the Gaussian standard deviation.

Quantification of the radial distributions (Figure S5A-F and Table S2) was performed on YZ transverse projections (obtained from 4×4×4 nm/ voxel, 800 nm thick YZ reconstructions from 3D-STORM images) using a custom ImageJ script. 160 nm wide line profiles were drawn across the axon submembrane (3 to 10 per transverse sections through apical, lateral and ventral AIS membranes), and intensity profiles for each channel were fitted with a Gaussian curve. Difference between the Gaussian means for each channel provided the distance between epitopes.

Quantification of the AIS position along the axon on epifluorescence images (Figure 6, S3 and S6) was also performed on the ßIV-spectrin SD labeling using a custom ImageJ script. A line profile (3.35 μm wide) was drawn along the axon, starting at the emergence point from the soma, and the intensity profile was smoothed using a 5 μm wide sliding window. The begin and end of the AIS for a given labeling were defined as the position along the smoothed profile where intensity first rose above 30% and last dropped below 30% of the maximum intensity value, respectively (Grubb and Burrone, 2010). Resulting AIS ROIs were used to measure mean intensities in other channels.

### Data visualization and statistics

Intensity profiles, graphs and statistical analyses were generated using Prism (GraphPad software). For all Figure legends, n refers to the total number of pooled individual measurement, and N to the number of independent experiments. For spacing graphs, the histogram of spacing values is overlaid with the corresponding Gaussian fit. On other graphs, dots are individual measurements, bars or horizontal lines represent the mean, and vertical lines are the SEM unless otherwise specified. Significances were tested using two-tailed unpaired t-tests (two conditions) or one-way ANOVA followed by Tukey post-test (3 or more conditions). For Gaussian fits of the spacing histograms, significance of the difference between the Gaussian means and standard deviations (Table S1) was determined using the fit error and the number of histogram bins + 1 as N (N=17). In all Figures and tables significance is coded as: ns non-significant, • p < 0.05, •• p < 0.01, ••• p < 0.001.

